# Temporal and spatial staging of lung alveolar regeneration is determined by the grainyhead transcription factor *Tfcp2l1*

**DOI:** 10.1101/2022.08.23.504977

**Authors:** Fabian L. Cardenas-Diaz, Derek C. Liberti, John P. Leach, Apoorva Babu, Jonathan Barasch, Tian Shen, Maria A. Diaz-Miranda, Su Zhou, Yun Ying, Michael P. Morley, Edward E. Morrisey

**Author notes:** Corresponding Author and Lead Contact: Edward E. Morrisey, Ph.D., University of Pennsylvania, Smilow Translational Research Center, Room 11-124, 3400 Civic Center Boulevard, Building 421, Philadelphia, PA 19104-5129.

## Abstract

Alveolar epithelial type 2 (AT2) cells harbor the facultative progenitor capacity in the lung alveolus to drive regeneration after lung injury. Using single cell transcriptomics, software-guided segmentation of tissue damage, and *in vivo* lineage tracing, we have identified the grainyhead transcription factor Tfcp2l1 as a key regulator of this regenerative process. Tfcp2l1 expression is initiated late in lung development and restricted to the AT2 cell population in the postnatal lung. Loss of Tfcp2l1 in adult AT2 cells decreased self-renewal and enhanced AT2-AT1 differentiation during active tissue regeneration. Conversely, Tfcp2l1 blunts the proliferative response to inflammatory signaling during the early acute phase after injury. This ability of Tfcp2l1 to temporally regulate the balance of AT2 self-renewal and differentiation is spatially restricted to zones undergoing active alveolar regeneration. Single-cell transcriptomics and lineage tracing reveal that Tfcp2l1 regulates cell fate dynamics by balancing the traffic across the AT2-AT1 differentiation axis and restricting the inflammatory program in AT2 cells. Organoid modeling shows that these cell fate dynamics are controlled by Tfcp2l1 regulation of IL-1 receptor expression and activity in AT2 cells. Together, these studies reveal the critical importance of properly staging lung alveolar regeneration and the integral role of Tfcp2l1 plays in balancing epithelial cell self-renewal and differentiation in this process.

## INTRODUCTION

Multiple forms of acute lung injury, including viral infections, can lead to acute respiratory distress syndrome (ARDS), exemplified by the COVID-19 pandemic (Torres Acosta and Singer, 2020). A critical challenge for the lung facing viral infection is to clear the pathogen while maintaining tissue function, particularly gas exchange within alveoli (Boyd et al., 2020; Flerlage et al., 2021; Torres Acosta and Singer, 2020). This delicate balance involves as yet poorly understood crosstalk between the immune system and alveolar cell lineages, including alveolar epithelial cells. The alveolar epithelium consists of two major cell types, alveolar epithelial type 1 (AT1) cells, which cover 95% of the externalized lumen of the alveolus and are critical for gas exchange, and alveolar epithelial type 2 (AT2) cells, which secrete pulmonary surfactant to prevent alveolar collapse during respiration. The AT2 cell population also harbors the facultative progenitor activity in the alveolar epithelium demonstrated by their ability to proliferate and differentiate into AT1 cells after injury and during homeostasis (Basil et al., 2020; Hogan et al., 2014; Liberti et al., 2021). Thus, understanding the transcriptional network regulating AT2 cell proliferation and differentiation is essential to developing new methods that promote lung regeneration.

The heterogeneous nature of lung damage and regeneration complicates the accurate assessment of alveolar injury models. Recent developments in single-cell technologies and computational tools for genomic analysis and the unbiased segmentation of tissue damage have helped to deconvolute this complexity and facilitate our understanding of the regenerative process in the lung (Choi et al., 2020; Kobayashi et al., 2020; Liberti et al., 2022; Strunz et al., 2020; Zepp et al., 2021). These tools have identified several molecular regulators of AT2 cell behavior during regeneration, including Wnt, Stat3, Fgf, and IL-1 signaling (Choi et al., 2020; Katsura et al., 2019; Liberti et al., 2021; Nabhan et al., 2018; Paris et al., 2020; Strunz et al., 2020; Zacharias et al., 2018; Zepp et al., 2021; Zepp et al., 2017). Despite this recent progress, the transcriptional networks that regulate the balance between AT2 self-renewal and differentiation during lung regeneration remain incompletely understood.

Transcription factor cellular promoter 2 like 1 (Tfcp2l1) is a member of the Grainyhead transcription factor family and is a downstream target of the Wnt and LIF/Stat3 signaling pathways (Hancock et al., 2021; Heo et al., 2020; Liu et al., 2017; Qiu et al., 2015). Tfcp2l1 is necessary and sufficient to maintain mouse embryonic stem cell (mESC) self-renewal and stemness and is also important for kidney development (Hancock et al., 2021; Heo et al., 2020; Sun et al., 2018; Wang et al., 2019; Werth et al., 2017; Zhang et al., 2021). Additionally, Tfcp2l1 regulates proliferation in multiple cancer cell types (Kotarba et al., 2018; Tun et al., 2010). Like other grainyhead transcription factors, Tfcp2l1 contains a CP2-like DNA binding domain and a dimerization motif through which it can homo- and heterodimerize with other grainyhead transcription factors (Taracha et al., 2018). However, little is known about whether Tfcp2l1 is expressed in or plays a role in lung regeneration.

In this study, we have identified Tfcp2l1 as a critical transcriptional regulator of AT2 self-renewal and AT2-AT1 cell differentiation in response to multiple models of acute lung injury. Using single-cell RNA-seq (scRNA-seq) and lineage tracing techniques, we show that Tfcp2l1 expression is initiated just before birth in the AT2 cell lineage and continues to be expressed throughout adulthood. Loss of Tfcp2l1 leads to decreased self-renewing capacity in the AT2 cell lineage with a simultaneous increase in AT2-AT1 cell differentiation in a spatiotemporal specific manner after acute lung injury. Mechanistically, we show that Tfcp2l1 regulates an AT2 cell inflammatory code that is important for their response to certain inflammatory cytokines, including IL-1β. Single-cell lineage tracing and transcriptomics reveal that the anchoring response of AT2 cells to acute injury changes upon loss of Tfcp2l1, leading to the primacy of the inflammatory expression program over cell proliferation. Our studies identify a new transcriptional regulator, Tfcp2l1, that is essential for organizing the stage-dependent response of AT2 cells to acute injury and their subsequent response during alveolar regeneration in the lung.

## RESULTS

### Tfcp2l1 expression is restricted to alveolar epithelial type 2 cells in the lung

To identify putative transcriptional regulators of AT2 cell progenitor function, we assessed gene expression differences between the Wnt responsive AT2 subpopulation called alveolar epithelial progenitors (AEP) and the entire AT2 cell population in the adult mouse using RNA-seq (Zacharias et al., 2018). We filtered this dataset to identify transcription factors with enhanced expression in AEPs, which revealed *Tfcp2l1* as one of the top differentially expressed genes **(Figure 1A)**. Analysis of adult lung single-cell RNA sequencing (scRNA-seq) data revealed that *Tfcp2l1* expression is restricted to AT2 cells in the lung (**Figure 1B-C)** (Zepp et al., 2021). Examination of a developmental time course scRNA-seq dataset revealed that *Tfcp2l1* transcript was detectable by scRNA-seq starting at approximately embryonic day (E) 17.5 (**Figure 1D**) (Zepp et al., 2021). To further validate this late onset of Tfcp2l1 gene expression, we used a *Tfcp2l1*^*CreERT2*^; *R26R*^*tdTomato*^ mouse line to track expression during development and into the adult. These data showed that Tfcp2l1+ cells could be identified by tamoxifen-induced recombination starting at E18.5, and expression was restricted to a subset of AT2 cells **(Figure 1E)**.

**Figure 1.**
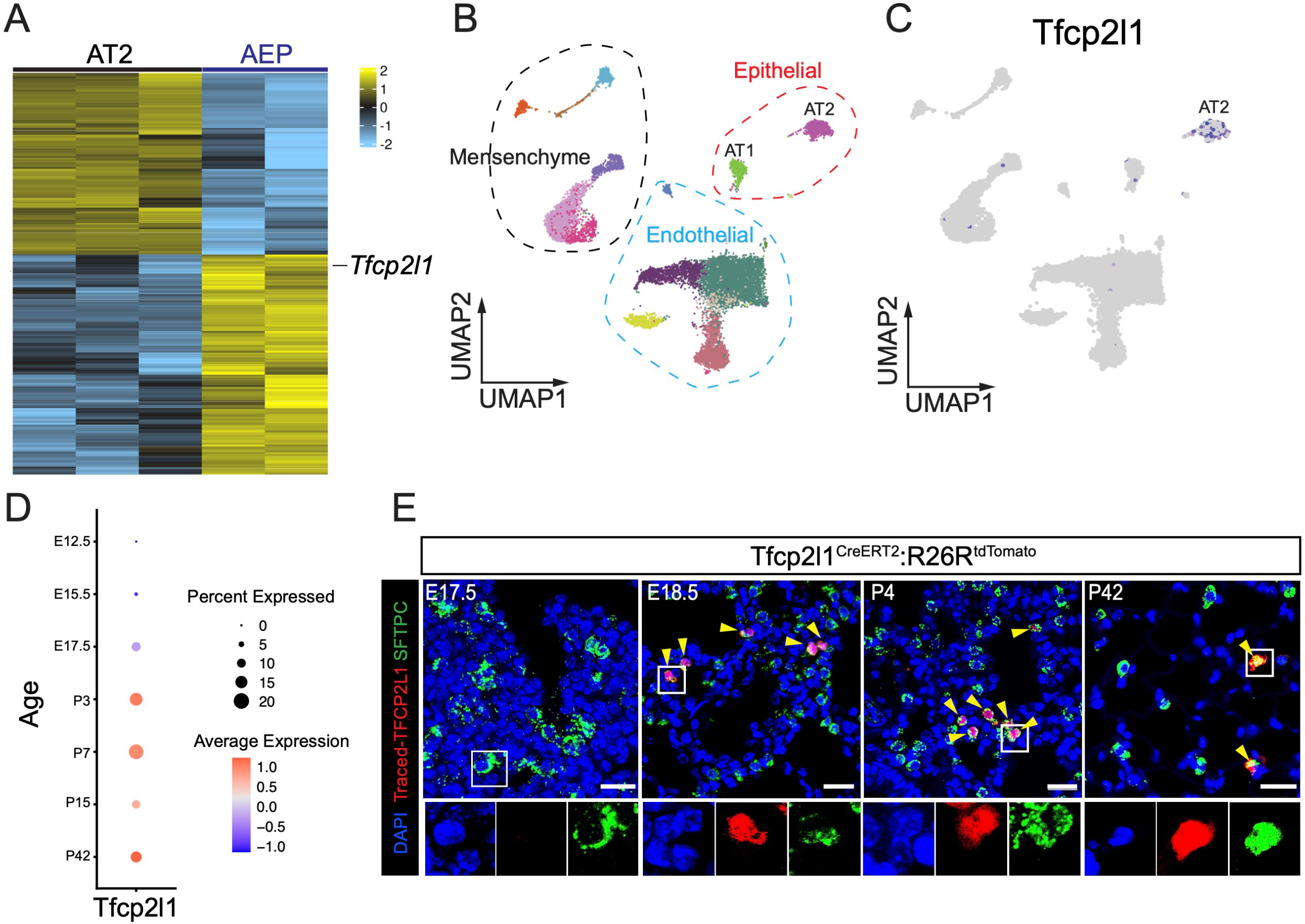
Tfcp2l1 expression is restricted to alveolar epithelial type 2 cells in the lung. (A) RNA sequencing heatmap plot showing differentially expressed genes between AT2 and AEP cell population. (B-C) Single-cell RNA sequencing UMAP plot of adult mouse lung with clustering and cell type distribution showing Tfcp2l1 expression in AT2 cells of the mouse lung. (D) DotPlot visualization derived from developmental time series of scRNA-seq data showing Tfcp2l1 expression during mouse lung development. (E) Time-specific lineage tracing using Tfcp2l1CreERT2:R26RtdTomato mice shows that Tfcp2l1 expression is initiated by approximately E18.5 and is restricted to Sftpc+ AT2 cells. Yellow arrows highlight lineage Tfcp2l1+/Sftpc+ cells. (Scale bar 20 µm).

### Loss of Tfcp2l1 in AT2 cells decreases self-renewal and increases AT1 cell differentiation during influenza-induced lung regeneration

To determine the role of Tfcp2l1 in adult AT2 cells, we generated *Sftpc*^*CreERT2*^; *Tfcp2l1*^*fl/fl*^; *R26R*^*EYFP*^ mice (from hereon called Tfcp2l1^AT2-KO^), allowing us to inactivate Tfcp2l1 in AT2 cells with a ∼90% efficiency (**Supplemental figure 1A-B**), and trace their fate in response to injury (**Figure 2A**). To assess the role Tfcp2l1 plays after an acute infectious injury to the lungs, we subjected Tfcp2l1^AT2-KO^ and control littermates to influenza A infection (H1N1), which results in severe and heterogeneous tissue damage followed by tissue regeneration (Katsura et al., 2019; Liberti et al., 2021; Liberti et al., 2022; Xi et al., 2017; Zacharias et al., 2018). During the acute inflammatory phase of influenza A infection (7-10 days post-infection), Tfcp2l1^AT2-KO^ mutants exhibited less weight loss than control mice suggesting a possible improvement in repair and regeneration (**Supplemental Figure 1C-E)**. This difference normalized as both Tfcp2l1^AT2-KO^ and control mice recovered. Lungs from Tfcp2l1^AT2-KO^ mutants and control mice were collected on day 14 post-viral infection (14 dpi) to examine differences in lung regeneration from loss of Tfcp2l1 expression. As previously reported, we segmented the regions of injury and regeneration into three distinct zones: normal, damaged, and severe (**Figure 2B-C)** (Liberti et al., 2021; Liberti et al., 2022). In normal zones, lung tissue exhibits largely homeostatic morphology with a typical distribution of AT2 cell (Sftpc+) and AT1 (Ager+) cells (**Figure 2C**). In severe zones, the tissue architecture is greatly altered, and there is an almost complete loss of alveolar epithelial cell markers with a few highly proliferative AT2 cells remaining (**Figure 2C-F**). However, in damaged zones that surround severe zones, we observe perturbed tissue morphology with dense accumulations of AT1 and AT2 cells (**Figure 2C)**. Examination of Ki67 immunostaining shows that Tfcp2l1^AT2-KO^ mutants exhibit less AT2 cell proliferation in damaged and severe zones (**Figure 2D-F)** (Liberti et al., 2021; Liberti et al., 2022). Assessment of cell proliferation at 28 days post-infection demonstrates that AT2 proliferation had decreased in both Tfcp2l1^AT2-KO^ mutants and controls with no significant difference between the genotypes, similar to homeostatic conditions (**Supplemental Figure 1, G-I**). Lineage tracing demonstrates that AT2-AT1 differentiation occurs at a much higher rate in Tfcp2l1^AT2-KO^ mutants in damaged zones at 14 dpi but normalizes by 28 dpi **(Figure 2G-I Supplemental Figure 1J-L**). These results suggest that Tfcp2l1 suppresses AT2 cell differentiation and promotes self-renewal after acute injury (**Figure 2J**).

**Figure 2.**
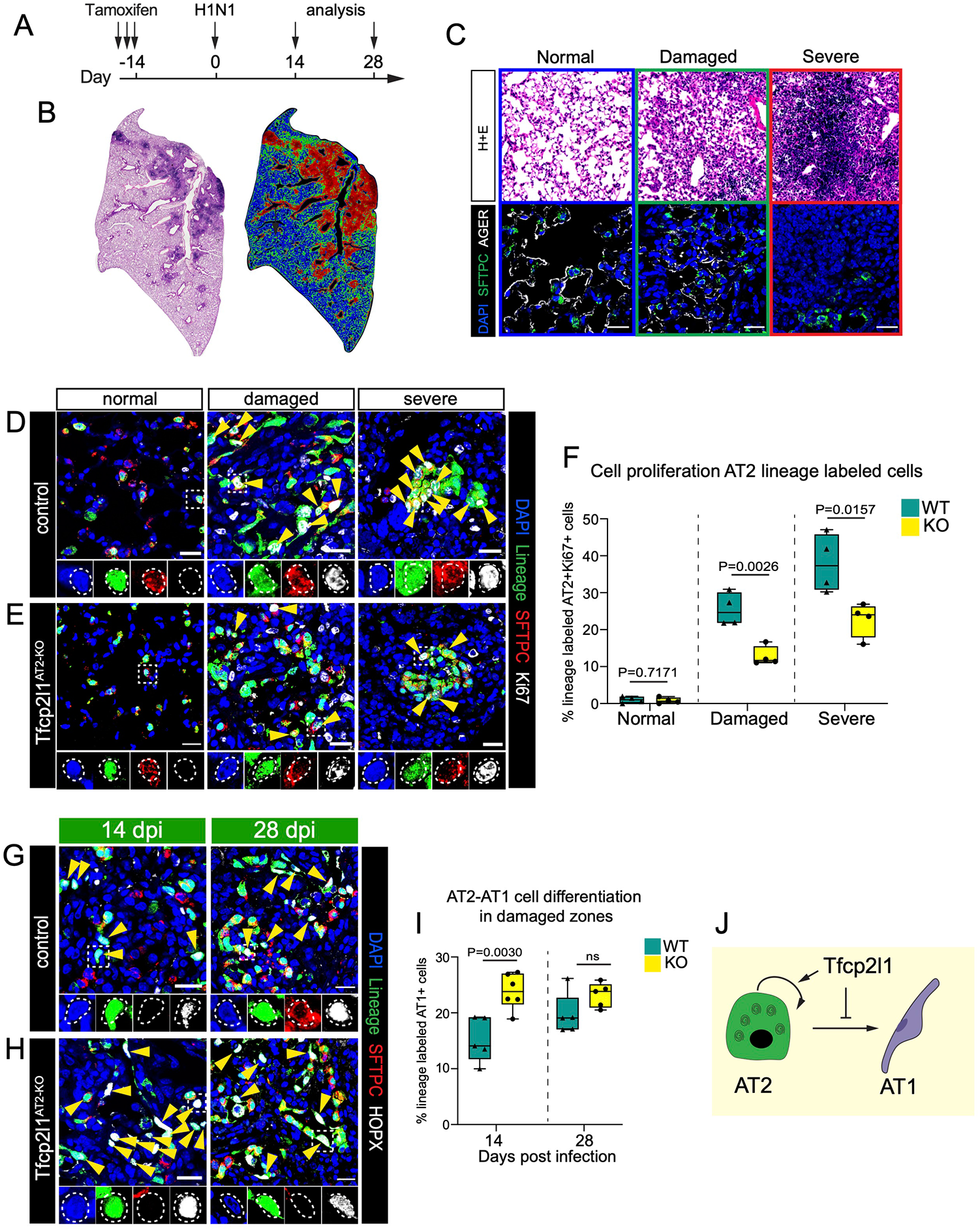
Loss of Tfcp2l1 in AT2 cells decreases self-renewal and increases AT1 cell differentiation during influenza-induced lung regeneration. (A) Experimental plan showing tamoxifen treatment and influenza infection and different timing for analysis. (B) Left: H+E stain at 14 days post-infection (dpi). Right: Cluster injury zone map generated from the H+E picture. (C) Top row: H+E pictures representing the injury zones found at 14 dpi. Color outlining each boxed represent different injury zones (blue=normal, green=damaged, red=severe; scale bar 100µm). Bottom row: IHC pictures for SFTPC, AGER, and DAPI in the three different injury zones (scale bar 20 µm). (D-E) IHC for the AT2 cell lineage marker EYFP, SFTPC, and Ki67 in normal, damaged, and severe injury zones at 14dpi with highlighted areas to show co-staining of markers (scale bar 20µm). (F) Quantification of lineage traced proliferative AT2 cells in different injury zones at 14 dpi. (G-H) IHC for the AT2 cell lineage marker EYFP, SFTPC, and HOPX in damaged zones at 14dpi (left) and 28dpi (right) with highlighted areas to show co-staining of markers and yellow arrow indicated AT1 cell-derived AT2 cells (scale bar 20µm). (I) Quantification of AT1 cell-derived AT2 cells in damaged injury zone at 14 and 28 dpi. (J) Summary diagram showing that Tfcp2l1 represses AT1 cell differentiation and promotes AT2 cell proliferation at 14dpi. All quantification data are represented as mean ± SEM. Two-tailed t tests p values shown, n= 4-5 mice per group.

### Tfcp2l1 regulates AT2 cell-mediated alveolar regeneration in a spatial and temporal manner

Our data show that Tfcp2l1 regulates the balance between proliferation and differentiation at 14 days post influenza infection. To determine if Tfcp2l1^AT2-KO^ cells have altered proliferation kinetics earlier in response to injury, we employed a non-infectious hyperoxia-induced acute lung injury (HALI) model to induce a more homogeneous lung injury (**Figure 3A**) (Amarelle et al., 2021; Liberti et al., 2022; Mach et al., 2011; Matute-Bello et al., 2008; Penkala et al., 2021). The HALI model allows for precise timing of injury and regeneration due to the lack of ongoing infection and damage in the infectious influenza model. To interrogate cell proliferation during the early inflammatory phase of this injury prior to AT1 cell differentiation, we subjected Tfcp2l1^AT2-KO^ mutants and controls to 72 hours of HALI. Mice were then pulsed with EdU four hours before collecting lung tissue three days post-HALI. Flow cytometry revealed an increase in EdU+ *Tfcp2l1*-deficient AT2 three days post-HALI (**Figure 3B-C**). To determine whether this increase in AT2 cell proliferation is due to the earlier time point assessed or to a different injury model, we performed histological analysis on control and mutant mice at seven days post-HALI. At this time point, we observed the stochastic emergence of fibrosis consistent with previous reports (Amarelle et al., 2021; Chen et al., 2007; Liberti et al., 2022). We used the same unbiased computational imaging approach as previously described to bin our analysis into three injury zones: normal (variable AT2 cell proliferation), damaged (highly proliferative AT2 cells and AT2-AT1 differentiation), and severe (ACTA2+ fibrotic foci with few AT2 and AT1 cells) (**Figure 3D-G**) (Liberti et al., 2021; Liberti et al., 2022). Quantification of AT2 cell proliferation demonstrated a statistically significant decrease in Tfcp2l1^AT2-KO^ mutant AT2 cell proliferation seven days after HALI (**Figure 3H-J**). We also found that Tfcp2l1^AT2-KO^ AT2 cells differentiated into AT1 cells at a higher rate compared to controls seven days after HALI, similar to what we observed 14 days after influenza-induced lung injury (**Figure 3K-M**). These data suggest that Tfcp2l1 regulates a precise temporal response to acute lung injury, repressing early AT2 cell proliferation, then subsequently becoming essential for AT2 cell proliferation and AT2-AT1 cell differentiation during the peak of the regenerative process.

**Figure 3.**
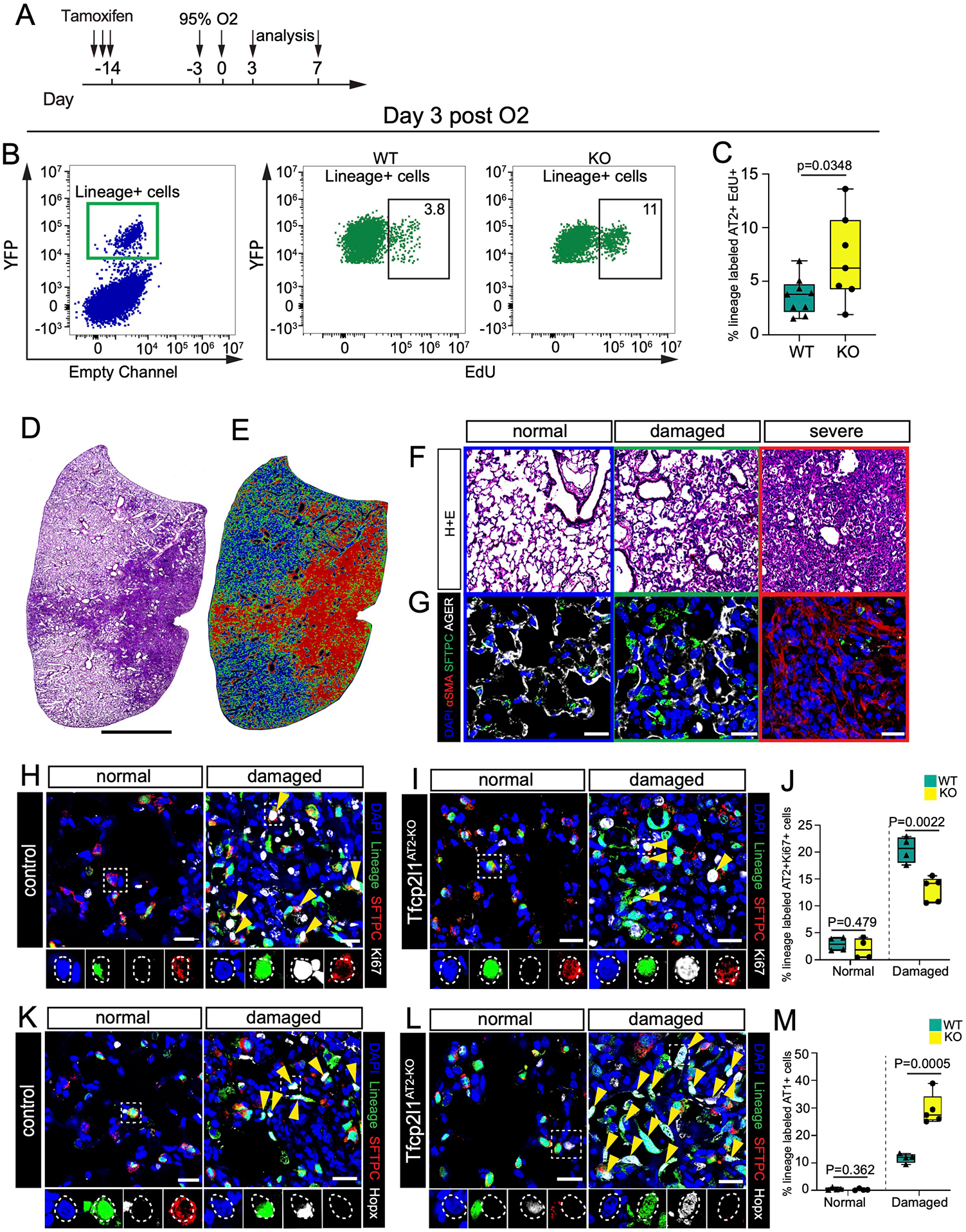
Tfcp2l1 regulates AT2 cell-mediated alveolar regeneration in a spatial and temporal manner. (A) Experimental schematic showing tamoxifen treatment and exposure to hyperoxia with different timing for analysis. (B) Flow cytometry quantification of EdU and lineage traced EYFP cells at day 3 post hyperoxia exposure. (C) Quantification of lineage traced EYFP and EdU positive cells at day 3 post hyperoxia exposure (n=7-9 mice per group). (D-E) Left: H&E stain at 7 days post hyperoxia exposure. Right: Cluster injury zone map generated from the H&E picture. (F) H+E pictures, each box panel is a representative picture of the injury zones found 7 days post hyperoxia exposure. Color in each boxed represent different injury zone (blue=normal, green=damaged, red=severe). (G) IHC pictures for SFTPC, AGER, and αSMA in the three different injury zones (scale bar 20 µm). (H-I) IHC for the AT2 cell lineage marker EYFP, SFTPC, and Ki67 in normal and damaged injury zones 7 days post hyperoxia exposure of control and Tfcp2l1AT2-KO mutants with dashed white boxes and yellow arrows highlighting proliferative lineage traced AT2 cells (scale bar 20µm). (J) Quantification of lineage traced proliferative AT2 cells in different injury zones at 7 days post hyperoxia exposure. (K-L) IHC for the AT2 cell lineage marker EYFP, SFTPC, and HOPX in normal and damaged zones 7 days post hyperoxia exposure of control and Tfcp2l1AT2-KO mutants with dashed white boxes and yellow arrows highlighting AT1 cells derived from AT2 cells (scale bar 20µm). (M) Quantification of AT1 cell-derived AT2 cells in normal and damaged injury zones at 7 days post hyperoxia exposure. All quantification data are represented as mean ± SEM. Two-tailed t-tests p values shown, n= 4-5 mice per group.

### Loss of Tfcp2l1 leads to altered AT2 cell transcriptional states and increased traffic across the AT2-AT1 differentiation axis

To determine how Tfcp2l1 regulates AT2-specific cell state changes during regeneration, we isolated Sftpc^CreERT2^ lineage traced cells from Tfcp2l1^AT2-KO^ and control animals 14 dpi by fluorescence-activated cell sorting (FACS) and performed single-cell RNA sequencing (scRNA-seq) analysis (**Figure 4A**). The 14-day post-influenza infection point was used since both proliferation and differentiation are occurring and affected by the loss of Tfcp2l1 at this time. Control and Tfcp2l1^AT2-KO^ mutant scRNA-seq datasets were merged, which revealed six distinct cell clusters (**Figure 4B-C**). As expected, we observed a cluster consisting of canonical AT1 cells, one cluster resembling the previously described transition state between AT2-AT1 cells (Choi et al., 2020; Kobayashi et al., 2020; Strunz et al., 2020), and four clusters correlating to various other subtypes of AT2 cells (**Figure 4C**). In addition, the AT2b and AT2/AT1 transition clusters were over-represented in the Tfcp2l1^AT2-KO^ mutants (**Figure 4D**). The AT2b cluster is marked by increased expression of genes relating to what was previously described as a primed or activated AT2 cell state, including *Lcn2, Dmkn*, and *Lrg1* (**Figure 4E**) (Choi et al., 2020; Strunz et al., 2020).

**Figure 4.**
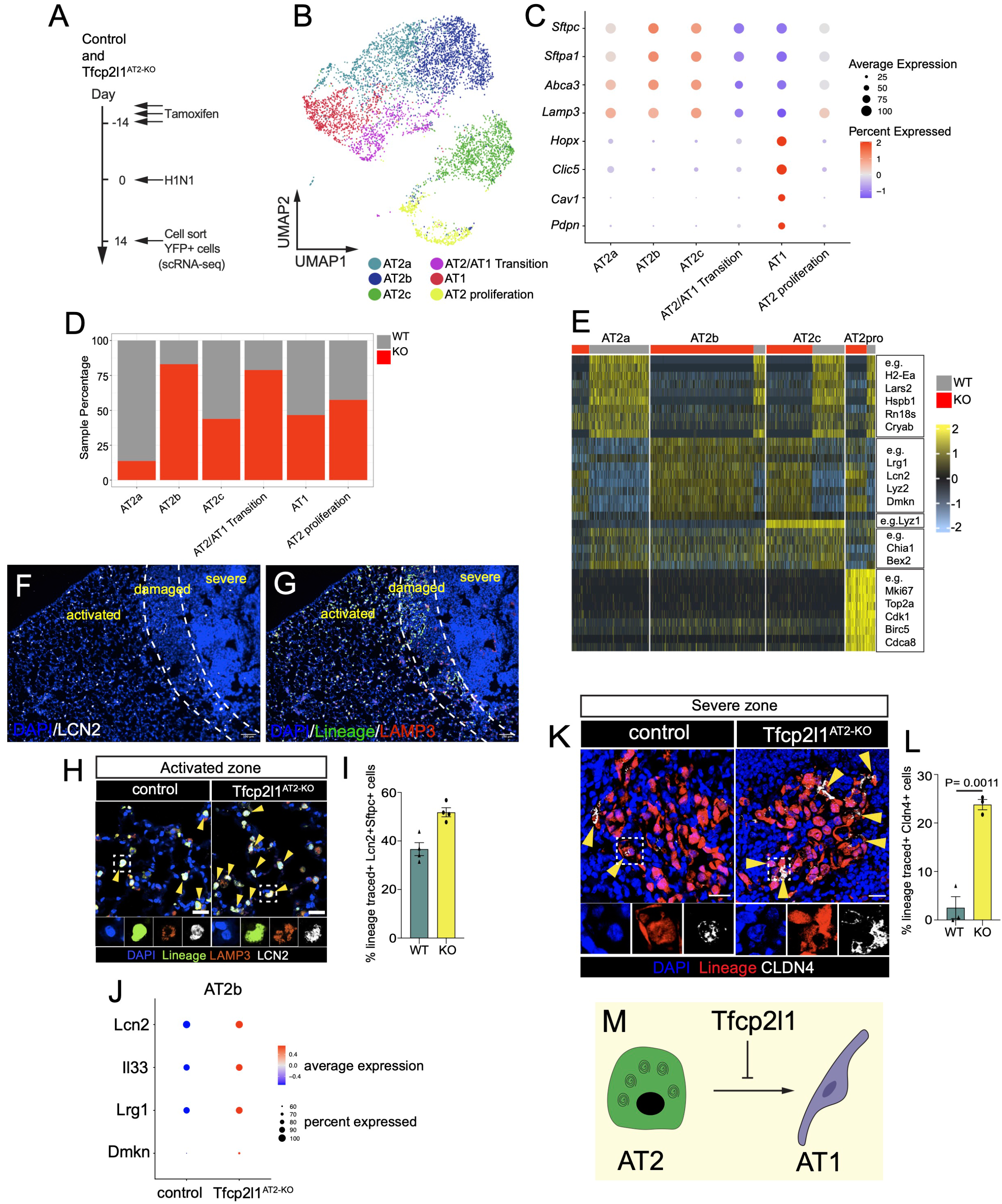
Loss of Tfcp2l1 leads to altered AT2 cell transcriptional states and increased traffic across the AT2-AT1 differentiation axis. (A) Experimental schematic showing tamoxifen treatment, influenza infection, and the timing to cell sort lineage traced EYFP cells to generate single-cell RNA sequencing (scRNA-seq) libraries from control and Tfcp2l1AT2-KO mice. (B) Merged scRNA-seq data, showing UMAP plot of control and Tfcp2l1AT2-KO mice 14dpi. (C) Dot plot graph from merged scRNA-seq data with AT1 and AT2 canonical markers by cell clusters. (D) Cell percentage distribution per cell cluster. (E) Heat map plot showing differential expressed genes per cell cluster. (F) IHC for LCN2 across the different zones of injury and regeneration at 14 dpi. (G) IHC for the AT2 cell lineage marker EYFP, LAMP3, and LCN2 in activated zones 14dpi. (H) IHC at high magnification of EYFP, LAMP3, and LCN2 in activated zones of control and Tfcp2l1AT2-KO mice at 14 dpi (scale bar 20µm). (I) Quantification of LCN2+ AT2 cells in activated injury zones at 14 dpi. (J). AT2b cell cluster dot plot displays Lcn2, Il33, Lrg1, and Dmkn expression between control and Tfcp2l1AT2-KO. (K) IHC for the AT2 cell lineage marker EYFP, and CLDN4 in severe zones at 14dpi, white boxes point at zoomed areas, and yellow arrow indicated CLDN4 positive cells in lineage traced cells (scale bar 20µm). (L) Quantification CLDN4+ lineage traced cells in severe injury zones at 14 dpi. (M) Summary diagram showing that Tfcp2l1 represses AT1 cell differentiation. All quantification data are represented as mean ± SEM. Two-tailed t-tests not significant; p≤ 0.05 n= 3-5 mice per group.

To determine which AT2 cell states emerge due to influenza infection and subsequent tissue regeneration, we compared uninjured AT2 cells to those exposed to influenza as described above at 14dpi. This comparison revealed three AT2-specific cell states after influenza injury, including one proliferative cluster (cluster 6) and three non-proliferative clusters (clusters 1, 2 and 4). (**Supplemental Figure 2A-B**). The major difference between the two non-proliferative clusters is the expression of the gene Lyz1 in cluster 2 versus cluster 1 (**Supplemental Figure 2D-F)**. Analysis of clusters 1 and 2 reveals the dramatic upregulation of pathways associated with immune effector response, antigen processing, and other immune-related phenomena (**Supplemental Figure 2G**). In addition, clusters 1 and 2 in this analysis are similar to the AT2b in the control versus Tfcp2l1^AT2-KO^ mutant analysis above (**Figure 4E**), indicating that loss of Tfcp2l1 leads to increased numbers of this inflammatory response state in AT2 cells (**Figure 4E and Supplemental Figure 2E and G)**.

We next assessed the location of AT2 cells expressing the inflammatory response pathways by examining Lcn2 expression, a marker of the inflammatory AT2b cluster, at 14 dpi in control and Tfcp2l1^AT2-KO^ mutant lungs (**Figure 4F**). Lcn2+ AT2 cells were present in control normal regions but found at lower numbers in damaged regions undergoing regeneration and essentially absent in the severe regions of both control and Tfcp2l1^AT2-KO^ mutants (**Figure 4F and G and Supplemental Figure 3A**). Interestingly, Lcn2+ AT2 cells were increased in number in Tfcp2l1^AT2-KO^ mutant normal zones (**Figure 4H-I)**. Moreover, the genes that define this inflammatory AT2b cluster are significantly increased in Tfcp2l1^AT2-KO^ mutant AT2 cells (**Figure 4J**). Since AT2 cells in what we previously described as a “normal” zone have elevated levels of an inflammatory gene signature, including Lcn2, we have renamed this region the “activated” zone to indicate that AT2 cells in morphologically unperturbed lung tissue react to injury and respond by activating an inflammatory program.

The AT2/AT1 transition cluster is enriched in genes previously identified as transitional AT2-AT1 cell state markers, including, *Ly6a, Tnip3*, and *Cldn4* (**Supplemental Figure 4B-C)** (Choi et al., 2020; Kobayashi et al., 2020; Strunz et al., 2020). To determine the relative abundance and localization of control and *Tfcp2l1*-deficient AT2-AT1 transition state cells during regeneration, we quantified the percentage of EYFP+ lineage traced cells expressing Cldn4 14 days after influenza infection in control and Tfcp2l1^AT2-KO^ mutants. While we observed only a small percentage of cells expressing Cldn4 in control animals, these cells were most frequently detected in severe zones and were increased in Tfcp2l1^AT2-KO^ mutants, suggesting increased traffic across the AT2-AT1 differentiation border (**Figure 4K-L**) **(Supplemental Fig 4D-E**). Thus, loss of Tfcp2l1 leads to alterations in the ratio of AT2 cell subsets, an increase in the AT2b inflammatory state, and increased traffic across the AT2/AT1 differentiation axis (**Figure 4M**).

### Loss of Tfcp2l1 disrupts stage specific transcriptional dynamics during AT2 cell regeneration

The alterations in the ratio of the inflammatory AT2b and AT2-AT1 transition state subpopulations suggested a change in the transcriptional response and trajectory of AT2 cells upon injury in the absence of Tfcp2l1. We applied the diffusion map (DM) algorithm to each scRNA-seq dataset as our data reduction approach. This method constructs potential differentiation trajectories but gives no directionality. We combined RNA velocity algorithms to predict direction in the DM (Haghverdi et al., 2015) **(Figure 5A-B)**. We next used scVelo to compute velocity vectors based on RNA expression and splicing parameters (**Figure 5C-D**) (Bergen et al., 2020; La Manno et al., 2018). While these methods can refine the directionally further by predicting cells age based on transcriptional dynamics referred to as “latent time “, a distinct advantage in our analysis is that we know the start state based on Sftpc^CreERT^ lineage tracing (homeostatic AT2 fate) and the end state (AT1 fate).

**Figure 5.**
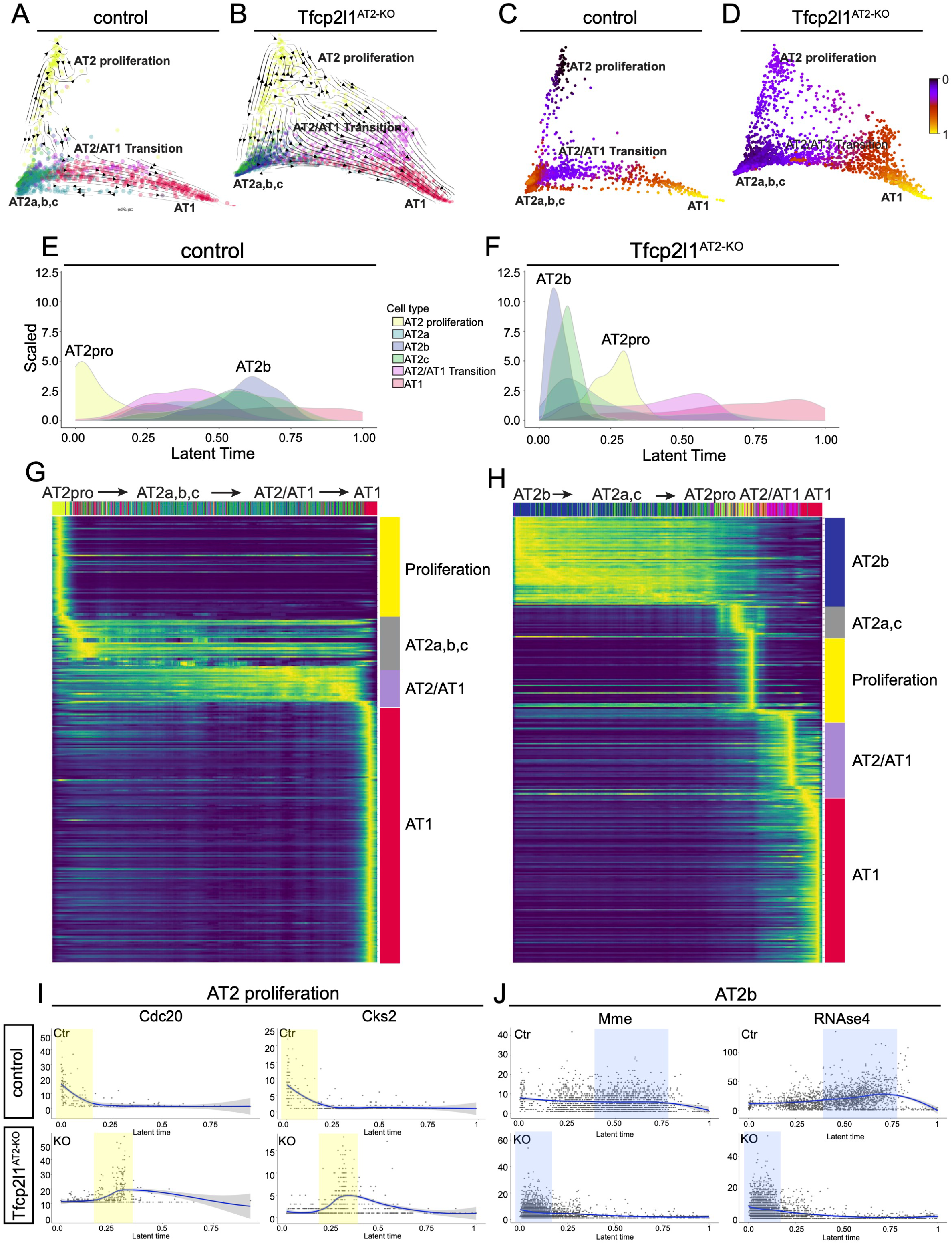
Loss of Tfcp2l1 disrupts stage-specific transcriptional dynamics during AT2 cell regeneration. (A-B) scVelo directionality overlaid onto diffusion map reduction of control and Tfcp2l1AT2-KO data at 14 dpi. (C-D) Latent time data inferred from scVelo mapped on the DM as a spectrum of transcriptional changes from a prime or 0 state (blue) to an end state (yellow) in control and Tfcp2l1AT2-KO mutants. Of note, both control and Tfcp2l1AT2-KO mutant trajectories end in the AT1 state. (E-F) Density histograms displaying the distribution of AT2 cell states based on latent time of lineage traced control and Tfcp2l1AT2-KO AT2 cells. (G-H) Gene expression dynamics resolved along latent time, showing AT2 cell state changes in lineage traced control and Tfcp2l1AT2-KO AT2 cells. (I-J) Expression dynamics of example driver genes along latent time (I) Cdc20 and Cks2, control (top), and Tfcp2l1AT2-KO (bottom). (J) Mme and RNAse4 control (top) and Tfcp2l1AT2-KO (bottom). Proliferation state shaded in yellow, AT2b state shaded in blue.

In control lungs, the AT2 cell proliferation state is the prime anchor state with AT1 cells being the final end state at 14dpi (**Figure 5A**). However, DM of Tfcp2l1^AT2-KO^ lineage traced AT2 cells shows that AT2a,b,c clusters are the prime anchor state with the proliferation state arising later in latent time (**Figure 5B**). This was confirmed by mapping latent time as a spectrum on top of the DM, which also shows a switch from proliferation to AT2a,b,c as the prime or 0 state for Tfcp2l1^AT2-KO^ lineage traced AT2 cells (**Figure 5C-D**) (Bergen et al., 2020; La Manno et al., 2018). To better understand the fate of AT2 cells upon loss of Tfcp2l1, linear ordering of the various cell states existing in the lineage traced cells was plotted using a histogram function and shows a clear switch of prime transcriptional states, with AT2b instead of AT2 proliferation positioned as the latent time origin in Tfcp2l1^AT2-KO^ traced cells (**Figure 5E and F**). In all analyses, both control and Tfcp2l1^AT2-KO^ mutant lineage traced cells end on the AT1 cell fate, which is supported by our lineage tracing (**Figure 5A-H**). These data suggest a reordering of the cell states that are derived from lineage traced Tfcp2l1 null AT2 cells 14 days after influenza infection, with a switch from the proliferative to inflammatory reactive AT2b state.

Genes which showed high degrees of transcriptional dynamics were used to assess the gene expression programs that differentially guided control and Tfcp2l1^AT2-KO^ mutant lineage traced AT2 cells through their response to acute injury (**Figure 5G and H**). Control lineage traced AT2 cells exhibited enrichment in cell growth, wound healing, and cell development gene expression programs (**Supplemental Fig. 5A**). In contrast, the response of Tfcp2l1^AT2-KO^ mutant lineage traced AT2 cells was driven by changes in cell adhesion, epithelial migration, and cell morphogenesis programs (**Supplemental Fig. 5B**). Latent time analysis shows that ordering of cells expressing two proliferation-associated genes, Cdc20 and Cks2, was altered between control and Tfcp2l1^AT2-KO^ mutants (**Figure 5I**). Similar but contrasting changes were observed by examining two AT2b specific genes, Mme and RNAse4 (**Figure 5J**). These data demonstrate that the loss of Tfcp2l1 disrupts AT2 cell states during lung regeneration leading to altered transcriptional dynamics, and a reordering of the staged response across latent time.

### Deep transcriptome analysis of lineage traced Tfcp2l1 deficient AT2 cells reveals an enhanced sensitivity to the post-injury inflammatory milieu

While scRNA-seq can provide important information regarding changes in cell state, the ability to quantitate changes in gene expression remains somewhat limited. To better understand the extensive quantitative changes in gene expression in AT2 cells as they respond to acute lung injury in the absence of Tfcp2l1 expression, we performed standard RNA-seq analysis on lineage traced cells from control and Tfcp2l1^AT2-KO^ mutants (**Figure 6A**). Differential gene expression between control and *Tfcp2l1*-deficient lineage traced cells revealed 386 genes upregulated and 195 genes down-regulated (**Figure 6B**). Gene set enrichment analysis identified several categories of genes down-regulated in Tfcp2l1^AT2-KO^ mutants, including those related to cell proliferation, such as E2F targets, MYC targets, and G2M checkpoint (**Figure 6C**). The top target genes driving the changes in the E2F proliferation-related pathway include *Cdc20, Cdkn3, Mxd3*, and *Top2a* (**Figure 6D**). Interestingly, genes related to inflammatory pathways were upregulated in *Tfcp2l1*-deficient lineage traced cells (**Figure 6C**). The top target genes driving upregulation of the inflammatory response pathway include *Il7r, Ccr7, Cd69, Il1a*, and *Il1r1* (**Figure 6E**). Additionally, we identified enrichment in AT1 cell markers in the *Tfcp2l1*-deficient cells, such as *Sema3e, Limch, Sema3a, Cav1, Clic5*, and *Hopx*, correlating with an increase in AT2-AT1 differentiation (**Supplemental Figure 5C)**. Several of these gene expression changes were confirmed by Q-PCR (**Figure 6F and G, Supplemental Figure 5D)**. These data reveal the dramatic changes in cell proliferation and inflammatory programs upon loss of Tfcp2l1 in regenerating AT2 cells.

**Figure 6.**
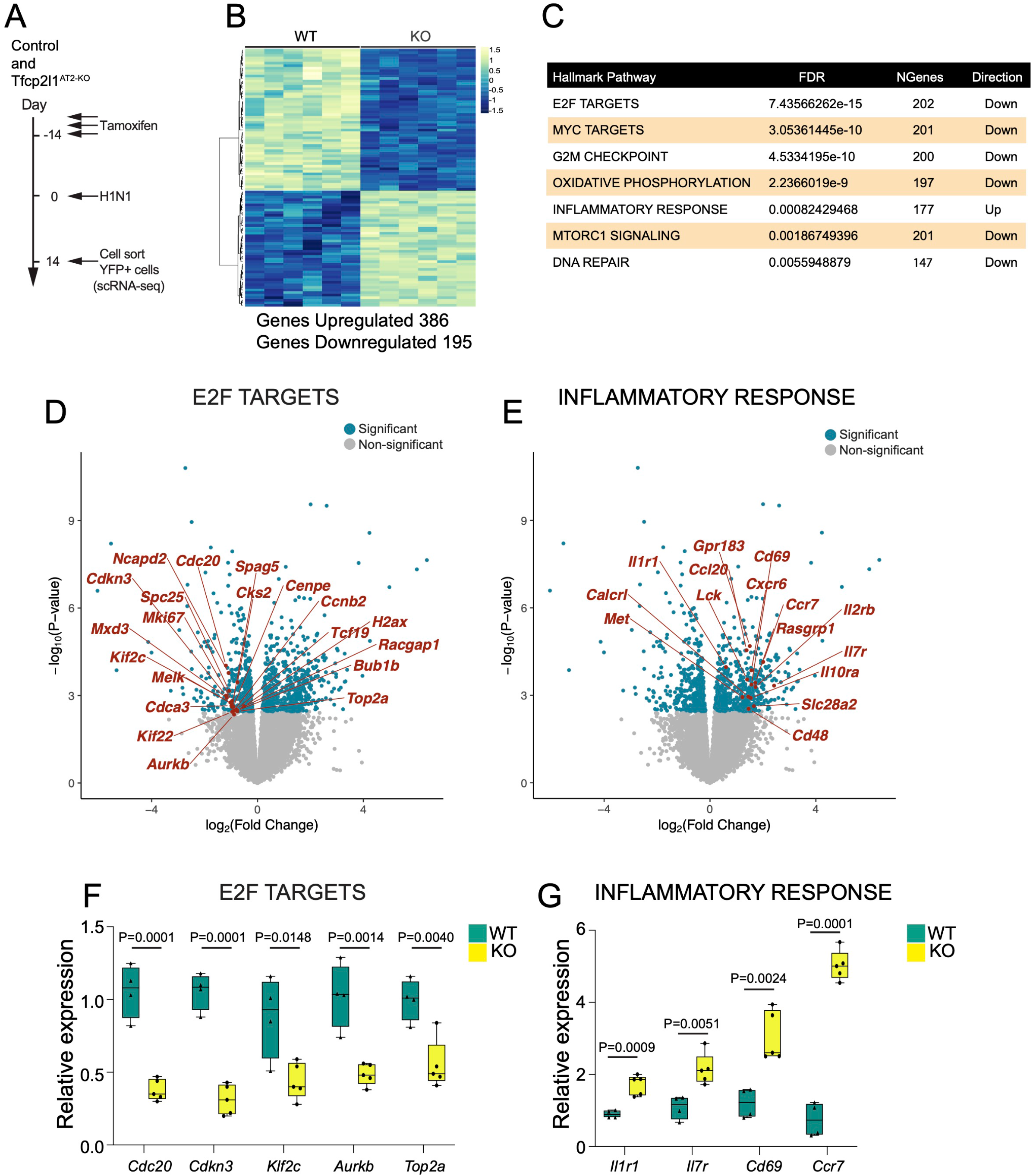
Deep transcriptome analysis of lineage traced Tfcp2l1 deficient AT2 cells reveals an enhanced sensitivity to the post-injury inflammatory milieu. (A) Experimental schematic showing tamoxifen treatment, influenza infection, and the timing to cell sort lineage traced EYFP cells to generate bulk RNA sequencing (RNA-seq) libraries from control and Tfcp2l1AT2-KO mice. (B) Differential expression heat map comparing control and Tfcp2l1AT2-KO mutants (n=6). (C) Hallmark database gene set enrichment analysis of differentially expressed genes between control and Tfcp2l1AT2-KO mice at 14 dpi. (D-E) Volcano plots from RNA-seq data Control and Tfcp2l1AT2-KO mice at 14 dpi, with dark blue dots showing statistically significant different genes between control and Tfcp2l1AT2-KO in the E2F and inflammatory response categories (adjusted p-value <0.05). Example gene in each category is highlighted in red. (F-G) Q-PCR gene expression validation of a subset of differentially expressed genes in the E2F and inflammatory response categories (Control n=4 and Tfcp2l1AT2-KO n=5).

### Tfcp2l1 restrains the AT2 cell response to IL-1 signaling

To better understand the response of Tfcp2l1-deficient AT2 cells to the inflammatory environment, we utilized an AT2 alveolar organoid model (**Figure 7A-B**)(Liberti et al., 2021; Liberti et al., 2022; Liberti and Morrisey, 2021). Since our transcriptomic analysis showed increased expression of multiple inflammatory cytokine receptors in Tfcp2l1-deficient AT2 cells including Il1r1 (**Figure 6E, G**), we examined whether these cells would respond in an enhanced manner to exogenous IL-1β exposure. IL-1 signaling regulates AT2 cell growth and proliferation (Choi et al., 2020; Katsura et al., 2019; Liberti et al., 2021; Liberti and Morrisey, 2021). While IL-1β treatment increased control organoid size, it led to an even greater increase in organoid size from Tfcp2l1-deficient AT2 cells (**Figure 7C**). These data suggest that Tfcp2l1 suppresses AT2 cell proliferation and AT2 cell responsiveness to IL-1β signaling (**Figure 7D**). Since inflammatory cytokine expression is highest in the early stages of post-injury repair and regeneration (Al-Garawi et al., 2009; Jang et al., 2012), these data support a model where Tfcp2l1 dampens the early response to the inflammatory milieu through control of Il1r1 and other cytokine receptors but is required in later stages of the regenerative process to balance AT2 self-renewal and AT2-AT1 differentiation (**Figure 7E**).

**Figure 7.**
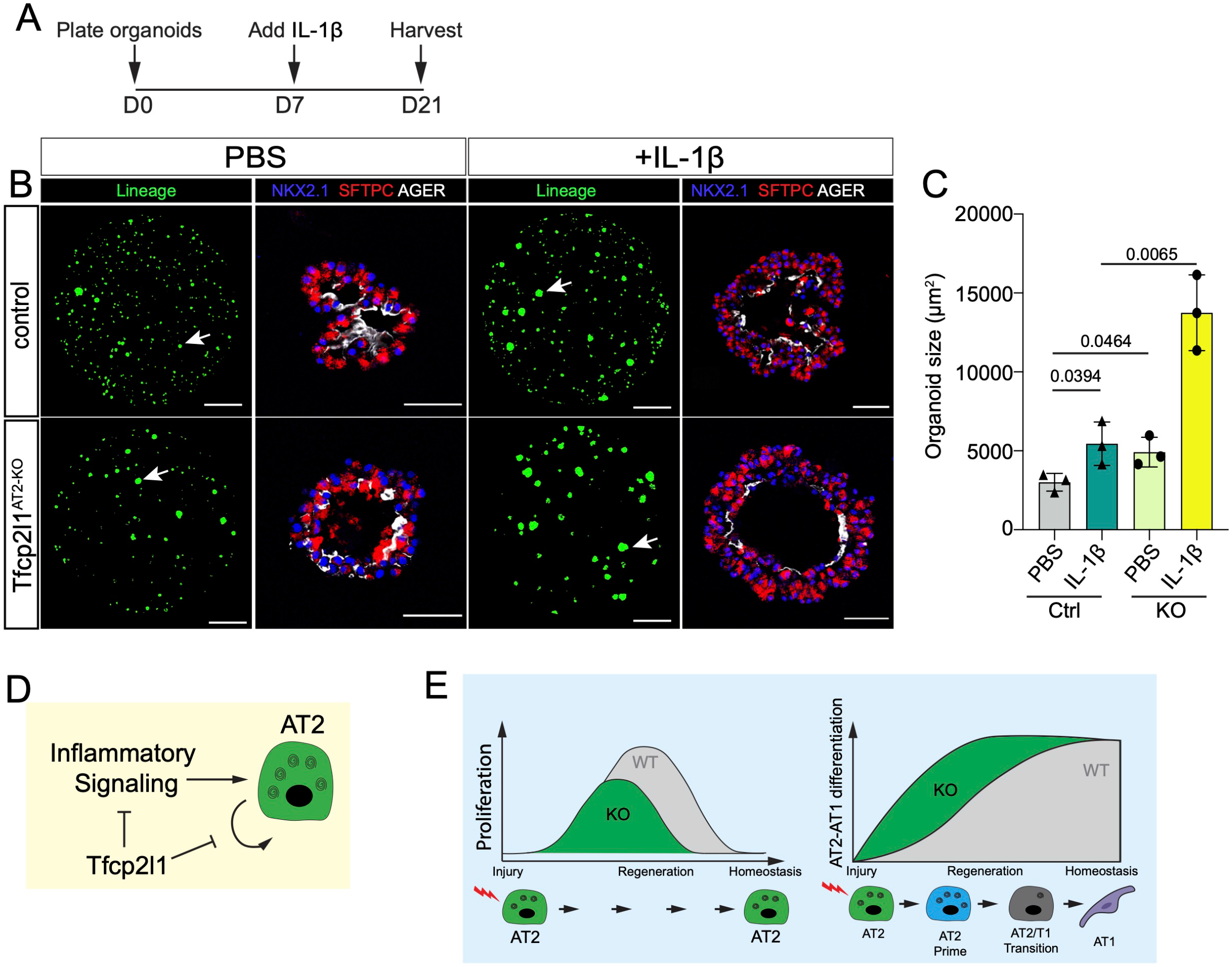
Tfcp2l1 restrains the AT2 cell response to IL-1 signaling. (A) Experimental schematic showing the days for cytokine treatments and duration of organoid culture. (B) Organoids treated with PBS or IL-1b as noted. Lineage-marked cells are from native EYFP fluorescence, and IHC on sections for SFTPC and AGER expression reveal AT2 and AT1 cells, respectively (lineage panel, bar graph 1000µm, IHC bar graph 50µm). (C) Quantification of organoid size after 21 days in culture comparing PBS and IL-1β in control and Tfcp2l1AT2-KO (D) Summary diagram showing that Tfcp2l1 represses AT2 cell proliferation and inflammatory signaling. (E) Summary diagram left: Tfcp2l1 is restricted to AT2 cells. The lack of Tfcp2l1 transient increases cell proliferation during lung regeneration. Right: Tfcp2l1 maintains AT2 cell identity and reduces AT2-AT1 cell differentiation. All data quantification is represented as mean ± SEM. Two-tailed t-tests not significant; p≤ 0.05 n=3 mice per group.

## DISCUSSION

The transcriptional networks regulating AT2 cell-mediated lung alveolar regeneration are not well understood. We identify Tfcp2l1 as an AT2 cell lineage-restricted transcription factor that regulates both AT2 cell self-renewal and differentiation in a spatial and temporal manner after acute injury. Tfcp2l1 regulates a complex transcriptional network that includes cell cycle drivers and inflammatory signaling, repressing the AT2 cell response to the exaggerated levels of inflammatory cytokines during the acute response early after injury, but promoting AT2 self-renewal and restricting AT1 differentiation later in the active regeneration process. Such precise balancing of the response by a resident epithelial progenitor is critical to ensure that it is not over-stimulated by the intense inflammatory environment observed after acute tissue injury to the lung but remains responsive to mitogenic signals later in the regenerative process.

Previous studies have shown that Wnt signaling drives and maintains AT2 cell fate during lung development and alveolar regeneration through the regulation of Wnt responsiveness in a subset of AT2 cells (Basil et al., 2022; Frank et al., 2016; Jacob et al., 2017; Liberti et al., 2021; Liberti et al., 2022; Nabhan et al., 2018; Zacharias et al., 2018). Tfcp2l1 is known to be a direct target of Wnt signaling, transmitting the actions of this pathway to regulate the pluripotent stem cell (PSC) state (Hancock et al., 2021; Liu et al., 2017; Qiu et al., 2015; Sun et al., 2018; Wang et al., 2019; Ye et al., 2013; Zhang et al., 2021). Our study shows that Tfcp2l1 expression emerges and increases in parallel with the second wave of Wnt signaling during sacculation and alveologenesis, which plays a key role in establishing and maturing the AT2 cell lineage (Frank et al., 2016). This expression timing and pattern also parallels the emergence of the Axin2+ AT2 subset called the alveolar epithelial progenitor (AEP) (Zacharias et al., 2018). The finding that loss of Tfcp2l1 leads to an increase in the rate of AT2-AT1 differentiation suggests that Wnt signaling, through Tfcp2l1, is an important but rate-limiting pathway balancing the epithelial regenerative response to acute injury. Such a balance is critical to respond rapidly after an acute injury to re-establish homeostasis, and Wnt responsive transcriptional regulators such as Tfcp2l1 play critical roles in balancing the proliferation-differentiation cycle.

AT2 cells are facultative stem cells within the lung alveolus, playing critical roles in surfactant production and maintenance and also exhibiting the capacity to rapidly re-enter the cell cycle and differentiate into AT1 cells (reviewed in (Leach and Morrisey, 2018). In PSCs, Tfcp2l1 is phosphorylated by Cdk1, which regulates its ability to promote cell proliferation (Heo et al., 2020). Tfcp2l1 also regulates Esrrb to promote PSC self-renewal (Wang et al., 2019). Increased expression of Tfcp2l1 in lung and bladder cancer has been linked to poor prognosis and aggressive phenotypes, whereas decreased expression has been linked to a decrease in cancer progression (Heo et al., 2020; Kotarba et al., 2018). Cdk1 phosphorylation of Tfcp2l1 also appears to regulate its ability to promote proliferation in bladder cancer (Heo et al., 2022; Heo et al., 2020). Together, these findings reveal an integral role for Tfcp2l1 in transmitting Wnt signals to promote cell cycle progression in multiple stem cell lineages, including AT2 cells.

Our original definition of the three major zones of responses to acute lung injury defined the morphologically unperturbed zone as “normal” (Liberti et al., 2021; Liberti et al., 2022). However, our current data reveals that AT2 cells exhibit a significant transcriptional response to acute lung injury in this region. Markers of the inflammatory reactive AT2b cluster such as Lcn2 are found almost exclusively in what we previously referred to as normal regions. This has led us to rename this region as “activated”. Activated AT2 cells in injured lungs increase overall, and their transcriptional response is enhanced upon loss of Tfcp2l1. While previous studies have identified similar AT2 cell subpopulations (Choi et al., 2020; Strunz et al., 2020), the spatial location of this unique subpopulation has remained unclear, as has the transcriptional network that regulates it. We show that *Tfcp2l1*-deficient AT2 cells upregulate IL1r1a, the receptor for IL-1β, and proliferate faster during early lung regeneration and in organoid cultures when exposed to exogenous IL-1β. These results suggest that Tfcp2l1 regulates AT2 cell receptivity to inflammatory signaling that occurs during the early stages of lung regeneration. However, later in the regenerative cycle, Tfcp2l1 is essential for balancing the proliferative and differentiation response of AT2 cells. One possible explanation for this bimodal response is that Tfcp2l1 is important in suppressing the proliferative response to an overly exuberant inflammatory setting where continuous mitogenic inputs into AT2 cells could increase their susceptibility to oncogenic conversion.

Several studies have identified the presence of a transient intermediate state during AT2-AT1 cell differentiation (Choi et al., 2020; Kobayashi et al., 2020; Strunz et al., 2020). At 14 days post influenza infection, we observed that this generally rare population is primarily restricted to severe zones where AT2 cells have not yet differentiated into AT1 cells. Loss of Tfcp2l1 in AT2 cells increases the number of cells expressing markers of this transient state in severe zones while increasing AT2-AT1 differentiation in the damaged zone. These results across different injury zones reinforce the concept that Tfcp2l1 maintains AT2 cell identity via multiple mechanisms, suppressing premature AT1 differentiation.

The molecular pathways that help balance the temporal response to acute tissue injury, particularly the altered inflammatory environment after injury, remain incompletely understood. Our studies suggest that Tfcp2l1 regulates both proliferative and inflammatory responsive gene programs to impede exuberant proliferation during the early inflammatory stage of acute injury but promote the necessary proliferation of AT2 cells later in the regenerative process after injury. Such tight control of these pathways underscores the necessity to control the epithelial response to acute injury and balance both the timing and extent of the pro-regenerative processes to achieve effective and functional tissue regeneration.

## AKNOWLEDGEMENTS

E.E.M is supported by funding from NIH grants (HL152194, HL087825, HL132999, HL134745) and the BREATH Consortium of Longfonds Foundation Netherlands. J.B is supported by funding from NIH grants (RO1DK073462, RO1DK092684, U54DK104309)

## AUTHOR CONTRIBUTION

F.L.C., D.C.L., J.P.L.; Performed experiments, F.L.C., D.C.L., J.P.L. E.E.M; design experiments, F.L.C., D.C.L., J.P.L., M.P.M., A. B., M.A.D., E.E.M.; Data acquisition and analysis, F.L.C., D.C.L., J.P.L., M.P.M., E.E.M.; Interpreted data, J.B., T.S., S.Z., Y.Y.; provided essential reagents and resources, F.L.C.; Draft writing, F.L.C., D.C.L., J.P.L., M.P.M., A. B., M.A.D., E.E.M.; Manuscript review and editing

## SUPPLEMENTARY DATA FIGURE LEGENDS

**Supplementary Figure 1**.

**(A)** Experimental schematic showing tamoxifen treatment to test Sftpc Cre-mediated Tfcp2l1 deletion in AT2 cells. **(B)** Relative gene expression analysis of Tfcp2l1 comparing control and Tfcp2l1AT2-KO mice. **(C)** Experimental schematic showing tamoxifen treatment and influenza infection, showing when mice weight was recorded. **(D)** Percentage of body weight for 14 days post influenza infection. **(E)** Body weight quantification from days 8 to 10. **(F)** Experimental schematic showing tamoxifen treatment, influenza infection, and the timing to examine mice lungs. **(G-H)** IHC for the AT2 cell lineage marker EYFP, SFTPC, Ki67 in normal and damaged zones at 28dpi, dashed white boxes point at zoomed areas, yellow arrow indicated proliferative lineage traced AT2 cells (scale bar 20µm) **(G)** Control, **(H)** Tfcp2l1AT2-KO. **(I)** Quantification of lineage traced proliferative AT2 cells in different injury zones at 28 dpi. **(J-K)** IHC for the AT2 cell lineage marker EYFP, SFTPC, and HOPX in Normal and severe zones at 14dpi (left) and normal zone 28dpi (right) dashed white boxes point at zoomed areas. Yellow arrow indicated AT1 cell-derived AT2 cells (scale bar 20µm), **(J)** Control, **(K)** Tfcp2l1AT2-KO **(L)** Quantification of AT1 cell-derived AT2 cells in damaged injury zone at 14 and 28 dpi.

**Supplementary Figure 2**.

**(A-B)** Merged scRNA-seq data, showing the UMAP plot of control AT2 cells isolated from mice not infected with influenza (No flu) compared to AT2 cells isolated 14dpi (Flu). **(A)** Cell cluster **(B)** Mouse AT2-no flu and AT2-flu sample distribution in different cell clusters. **(C)** Cell percentage distribution per cell cluster. **(D)** Feature plots showing distribution and expression of Lamp3, Hopx, Lyz1, Lcn2, and Lrg1 in mouse AT2-no flu and AT2-flu merged at 14dpi. **(E)** Heat map showing differential expressed genes per cell cluster comparing AT2-no flu and AT2-flu. **(F)** Dot plot visualization showing unique gene expression patterns in clusters 1, 2, 4, and 6. **(G)** GO analysis of clusters one and two.

**Supplementary Figure 3**.

**(A)** IHC for the AT2 cell lineage marker EYFP, SFTPC, and LCN2 in activated, damaged and severe zones 14dpi, white boxes point at zoomed areas, yellow arrow indicated LCN2 positive cells in AT2 cells (scale bar 20µm).

**Supplementary Figure 4**.

**(A)** Merged scRNA-seq data, showing UMAP plot of control and Tfcp2l1AT2-KO mice 14dpi. **(B)** Dot plot visualization showing unique gene expression pattern in merged AT2 control and Tfcp2l1AT2-KO mice 14dpi. **(C)** Top. Feature plots showing distribution and expression of Ly6a, Tnip3, and Cldn4. Bottom. Gene expression of Ly6a, Tnip3, and Cldn4 per cell cluster showed using violin plots in merged AT2 control and Tfcp2l1AT2-KO mice 14dpi. **(D)** IHC for the AT2 cell lineage marker EYFP and CLDN4 in activated, damaged severe zones at 14dpi, white boxes point at zoomed areas, yellow arrow indicated CLDN4 positive cells in lineage traced cells (scale bar 20µm). **(E)** Quantification CLDN4+ lineage traced cells is activated, damaged severe zones at 14dpi. All quantification data are represented as mean ± SEM. Two-tailed t-tests not significant; p≤ 0.05 n= 3 mice per group.

**Supplementary Figure 5**.

**(A-B)** GO enrichment analysis from putative genes in latent time analysis **(A)** Control **(B)** Tfcp2l1AT2-KO. **(C)** AT1 cell markers represented in volcano plot from RNA-seq data showing control and Tfcp2l1AT2-KO mice at 14 dpi (adjusted p-value <0.05) dark grey dots show statistically significant different genes between control and Tfcp2l1AT2-KO **(D)** Relative gene expression validation for AT1 cell target genes.

### Mouse Lines

Information related with mouse genotyping and mouse generation can be found from Jackson Laboratories otherwise is indicated. R26R^*EYFP*^ (Jackson Laboratory stock # 007903), R26R^*tdTomato*^ (Jackson Laboratory stock # 007914), Tfcp2l1^*CreERT2*^ (Jackson Laboratories stock # 028732). The Sftpc^*CreERT2*^ mouse line was a kind gift from Dr. Harold A. Chapman’s lab (University of California, San Francisco), and the Tfcp2l1^*flox*^ mouse was a kind gift from Dr. Barash Jonathon’s lab Columbia University). All mice were mixed background (C57BL/6 and CD1). We performed the experiments with at least n=3 mice per condition; we included mixed gender and carefully included littermates to ensure consistency. A single dot in each graph represents an individual mouse for data quantification. The University of Pennsylvania Institutional Animal Care and Use Committee approved all animal procedures performed here.

### Lung Alveolar organoid assay

Three consecutive tamoxifen doses (200mg/kg) were administered by oral gavage in Sftpc^*CreERT2*^; R26R^*E*YFP^ or Sftpc^*CreERT2*^; Tfcp2l1^*fl/fl*^; R26R^*EGFP*^ mice. 14 days later, mouse lungs were collected to generate single-cell suspension using a combination of different enzymes (Collagenase type I, dispase, and DNase1) as described in (Liberti et al., 2021; Liberti et al., 2022; Penkala et al., 2021). Samples were resuspended in FACS buffer (1X Phosphate-Buffered Saline (PBS), 25mM HEPES (Gibco, catalog # 15630080), 2mM EDTA (Invitrogen, catalog # 15575020), and 1.5% Fetal Bovine Serum (FBS) (ThermoFisher, catalog # 15260037). EGFP+ cells were FACS sorted, and we measured cell viability using Trypan Blue solution (Gibco, catalog # 15250061). To assess cell numbers, we used Countess 3 automated cell counter.

To generate alveolar organoids, we used a 24 well-plate format. The conditions described as follows used per one well. Each well consists of a cell culture inset (0.4µm transparent PET membrane) (Corning, cat # 353095). We mixed 5×103 *EGFP*+ cells with 5×104 primary lung fibroblasts to support epithelial cell growth with 50µl of ice cold Matrigel (Corning, catalog # 356231) and 50µl of MTEC-SAGM medium mixed with supplements (Lonza, catalog # CC-3119) and (Lonza, catalog # CC-4124), the SAGM medium preparation is detailed in (Liberti et al., 2022). The 100µl solution (Cells, medium, and Matrigel) was plated on top of the cell culture insert and let to polymerize for 20min in a tissue culture incubator (5%CO2, 37°C, with saturating humidity). Each well was fed using SAGM medium with a 10mM rock inhibitor (Y-27632 dihydrochloride, Millipore Sigma, catalog # Y0503) for 24 hours. Then, the medium was changed with fresh SAGM medium supplements with SB431542 (10µM, Tocris, Cat # 1614) for seven days changing the medium every other day. On day 7, organoids were fed using SAGM medium supplemented with or without IL-1β (10ng/mL, BioLegend, cat # 575102) until day 21, provided every other day.

We used EVOS FL Auto 2 Imaging System to document the organoid formation. For data analysis, we harvested the organoids on day 21. To perform IHC in organoids. First, organoids were fixed in the cell culture insert using 2% paraformaldehyde solution (Thermo Fisher Scientific, catalog # AAJ19943K2) for 30 minutes. Then, remove PFA using 3-5 washes in PBS, each for 10 minutes. After removing PFA, we proceeded to dehydrate organoids utilizing a series of ethanol washes from 30%, 50%, 70%, 95%, and 100% each wash for 30 minutes. Finally, the bottom of the cell culture inset was removed and embedded in paraffin wax, sectioned at a thickness of 6mm. Results are from three independent experiments, and each experimental condition was performed with at least three replicate wells.

### Tamoxifen delivery

Tamoxifen (Sigma-Aldrich, catalog # T5648) was resuspended at 20mg/mL using a mixture of ethanol (10%) and corn oil (90%). For pregnant dam experiments, tamoxifen was administered only once by oral gavage at a 200mg/kg dose. Experiments involve adult mice (older than six weeks old). We administer tamoxifen by oral gavage at a dose of 200mg/kg for three consecutive days. We left tamoxifen to wash out for 14 days for every injury experiment before starting injury experiments.

### EdU incorporation

EdU (Santa Cruz, catalog # sc-284628B) was dissolved in 1X PBS and administered via intraperitoneal injection at 50mg/kg. Adult mice received EdU 4 hours before lung collection. To detect EdU+ cells, we used the Click-iT EdU cell proliferation kit (Invitrogen, catalog # C10634) and followed the manufacturer’s instructions.

### Influenza infection

We administered intranasal 50µl of virus solution (PR8 H1N1 influenza virus in cold PBS) per anesthetized mouse. The viral dose is approximately 1LD50 (determined experimentally based on delivery to six-to eight-week-old C57BL/6 female mice) as previously described (Liberti et al., 2021; Liberti et al., 2022). We received the PR8 H1N1 influenza virus as a kind gift from Dr. John Wherry at the University of Pennsylvania.

### Histology

The lungs were harvested, and we removed blood from lung lobes using PBS by puncturing the heart and running PBS for one minute at a constant pressure of 25cm. Then we inflated the lungs using 2% PFA at continuous pressure of 25cm and left them fixed overnight at 4oC. To remove PFA, we wash the fixed lungs six times using PBS, every wash for 30 minutes, then proceed to dehydrate using 75%, 90%, and 100% ethanol each incubation for 24 hours at 4 oC. Lungs were paraffin wax, sectioned at a thickness of 6mm. Lung sections were stained using Hematoxylin and eosin (H&E) as performed before (Liberti et al., 2021; Liberti et al., 2022). To perform immunohistochemistry, we used the antibodies listed below using the following concentrations:

- HOPX (mouse, Santa Cruz, sc-398703, 1:100)
- Rabbit anti-SFTPC (Rabbit, Abcam, ab90716)
- ACTA2 (mouse, Millipore Sigma, A5228, 1:200)
- AGER (Rage) (rat, R&D Systems, MAB1179, 1:50)
- LAMP3 (DC-Lamp) (rat, Novus, DDX0191P-100, 1:100)
- GFP (chicken, Aves Labs, GFP-1020, 1:200)
- Ki67 (mouse, BD Biosciences, 550609, 1:200)
- NKX2.1 (Ttf1) (mouse, Thermo Fisher Scientific, MS-699-P1, 1:25)
- RFP (Rabbit anti-RFP, Rockland, 600-401-379)
- CLDN4 (Rabbit anti-Cldn4, ThemoFisher, 36-4800)
- LCN2 (Rabbit anti-Lcn2, Cell signaling, Cat No. D4M8L)

Slides were mounted using Vectashield Antifade Mounting Medium (Vector Laboratories catalog # H-1000) or Slowfade Diamond Antifade Mountant (Invitrogen, catalog # S36972). Slowfade Diamond Antifade Mountant was used to avoid the quenching action of Vectashield on Alexa Fluor 647 secondary antibodies. Imaging for cell quantification was acquired using a Leica TCS SP8 confocal microscope.

### Hyperoxia treatment and lung assessment program

We used adult mice (>6 weeks old) for hyperoxia experiments, and the methodology used was previously described. (Liberti et al., 2022; Penkala et al., 2021). In addition, we employed the lung assessment program to identify lung injury zones caused by influenza or hyperoxia injury to determine lung injury zones, and the software details are described (Liberti *et al*., 2022)

### Bulk RNA-seq and scRNA-seq

14 days post-infection lineage traced EGFP+ cells were FACS, and cell pellets were resuspended with lysis buffer from RNA isolation kit PureLink RNA Micro Kit (Invitrogen Cat No 12183-016). To isolate total RNA, we follow the manufacturing protocol. To calculate RNA integrity, we used a Bioanalyzer High Sensitivity RNA pico kit Analysis (Agilent, cat # 5067-1513). The NEBNext Single Cell/Low Input RNA Library Prep Kit for Illumina (New England Biolabs, catalog # E6420) follows the manufacturer’s instructions. Libraries were sequenced using the Illumina HiSeq. Fastq files were aligned against mouse reference (mm39/mGRC39) using the STAR aligner (v2.7.9a) (Dobin et al., 2013). Duplicate reads were removed using MarkDuplicates from Picard tools, and per gene read counts for Ensembl (v104) gene annotations were computed. Expression levels in counts per million (CPM) were normalized and transformed using VOOM in the limma R package (Ritchie et al., 2015). Surrogate variables to account for sources of latent variation such as batch were calculated using the svaseq function from the R SVA package (Leek et al., 2012). Differential gene expression analysis was conducted using the limma package. Gene Ontology and pathway analysis was performed using the clusterProfiler (v4.4.4) (Wu et al., 2021b). All plots were constructed in R using ggplot2 or Complexheatmap.

For scRNA-seq libraries reads were aligned to the mouse reference (mm39/mGRC39) and unique molecular identifier (UMI) counts obtained STAR-Solo (v2.7.9a). (Dobin et al., 2013). For further processing, integration and downstream analysis, Seurat (v4.0.6) (Hao et al., 2021) was used. Cells with less than 200 genes, greater than 2 Median absolute deviation above the median, and with potential stress signals of greater than 5% mitochondrial reads were removed. The cell cycle phase prediction score was calculated using Seurat function CellCycleScoring. Multiple libraries were merged and then merged data was normalized and scaled using the SCTransform function and regressing out the effects of percent fraction of mitochondria, number of features per cell, and number of UMI per cell. Linear dimension reduction was done via PCA, and the number of PCA dimensions was evaluated and selected based on assessment of an ElbowPlot. Data was clustered using the Louvain graph-based algorithm in R and Cluster resolution chosen based on evaluation by the clustree program. The Uniform Manifold Projection (UMAP) data reduction algorithm was used to project the cells onto two dimensional coordinates. Clusters were then assigned putative cell types based on annotation with canonical marker genes, or from assessment of top cluster-defining genes based on differential expression (using the FindAllMarkers function in Seurat). For intra-cluster gene expression differences, the FindMarkers function was used to identify variation between specified clusters and the resultant gene sets were comparted via the MAST method. GO and WikiPathway enrichment analysis was done with the clusterProfiler R package (Wu et al., 2021a). RNA velocity analysis was performed using the scVelo package. Count data for spliced, unspliced and ambiguous reads was obtained using velocity parameter in STAR-solo (v2.7.9a) and counts were converted to loom files using loompy. were created using scanpy (v1.9) (Wolf et al., 2018) and scVelo (v0.2.4) (La Manno et al., 2018) was used to compute RNA velocity and latent time. RNA velocity and heatmaps were generated using scVelo.

### Organoid size and colony forming efficiency

To measure organoid size, we follow the methodology described by (Liberti et al., 2021; Liberti et al., 2022). In brief, EVOS FL Auto 2 Imaging System was used to image the whole well to visualize the endogenous EYFP signal. We used Fiji software to process image analysis, every picture threshold, and further binarized. To minimize background, we applied erosion and then dilation. To fully measure organoids diameter, we applied to fill holes and watershedding functions. Finally, analyze particle was used to measure organoid size with reading parameter bigger than 1000^mm2.^ We performed at least three biological experiments with a minimum of 4 replicates.

### Flow cytometry

Samples were dispersed into single cells and then fixed using 2% PFA incubating at room temperature for 30min. Cells used for flow cytometry to detect EdU+ samples were processed using the Click-iT EdU cell proliferation kit (Invitrogen, catalog # C10634), and it was used according to the manufacturer’s instructions. Samples were analyzed using a CytoFLEX S flow cytometer.

### RNA isolation cDNA synthesis and Q-PCR

Total RNA was isolated using PureLink RNA Micro Kit (Invitrogen Cat # 12183-016). We resuspended the RNA in 16μl of RNAse-free, and we used 100ng total RNA to generate cDNA using the SuperScript™ IV Reverse Transcriptase (ThermoFisher Cat No 18090050). For Q-PCR, we mixed cDNA with SYBRgreen PCR master Mix (ThermoFisher Cat # 4367659). QuantStudio7 Flex (Applied Biosystems) was used for sample quantification. We used the delta-delta Ct method for gene expression quantification using the Tbp housekeeping gene for normalization. The different gene primers are presented in Table 1.

**Table 1.**
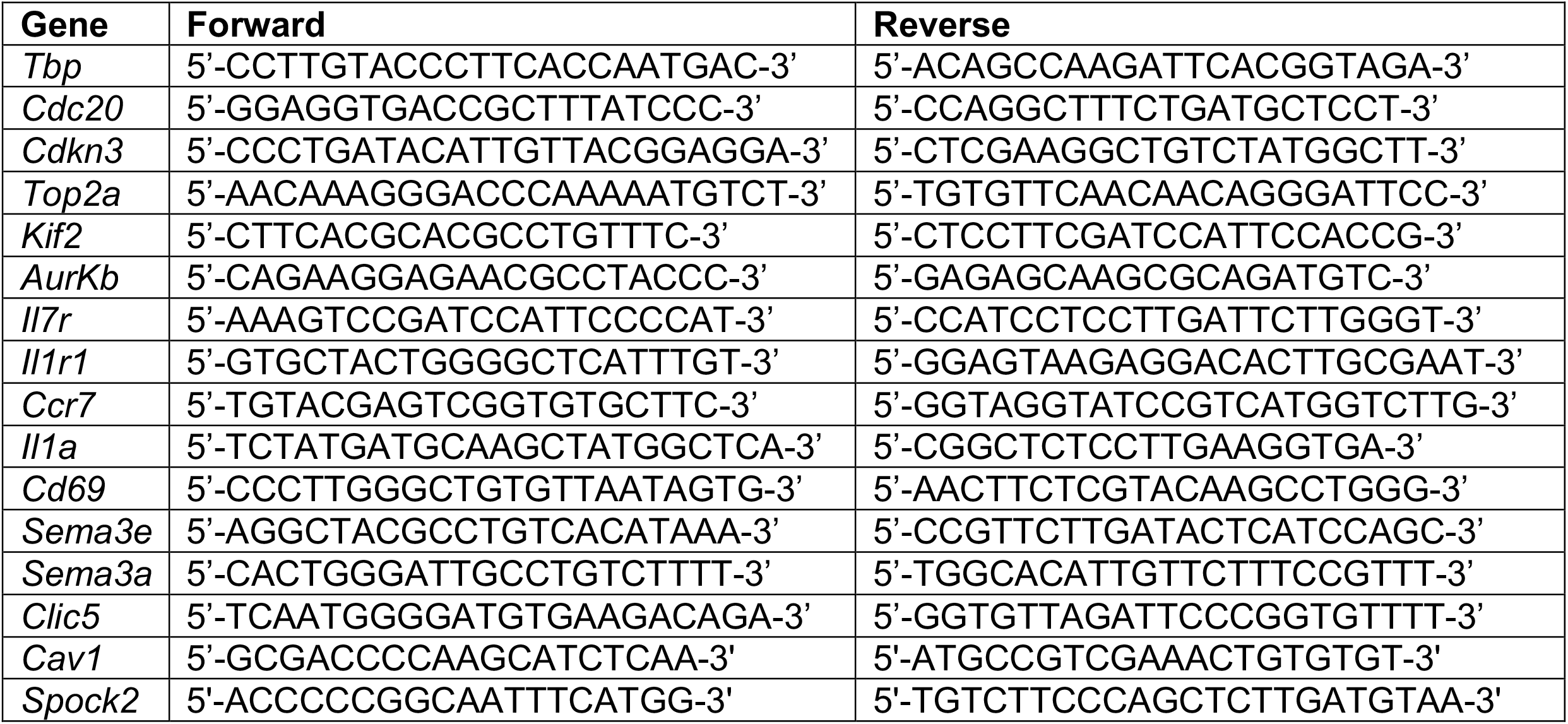
Q-PCR Primers.

### Quantification and Statistics

Results related to quantification based on images or any other type of experiment it is shown as mean ± SEM if another representation is not stated. To compare two different populations, we used two-tailed unpaired Student’s t-tests with p-values <0.05 considered significant. We used Graphpad Prism9 software to compute and graph data.

## DATA AND SOFTWARE AVAILABILITY

Tfcp2l1^AT2-KO^ and control RNA-seq and scRNA-seq datasets are deposited in the NCBI GEO data base at https://www.ncbi.nlm.nih.gov/geo/query/acc.cgi under GEO accession number GSE204787

