## Supplemental figures for "Temporal and spatial staging of lung alveolar regeneration is determined by the grainyhead transcription factor *Tfcp2l1*"

**Supplemental Figure 1. Cardenas et. al.**

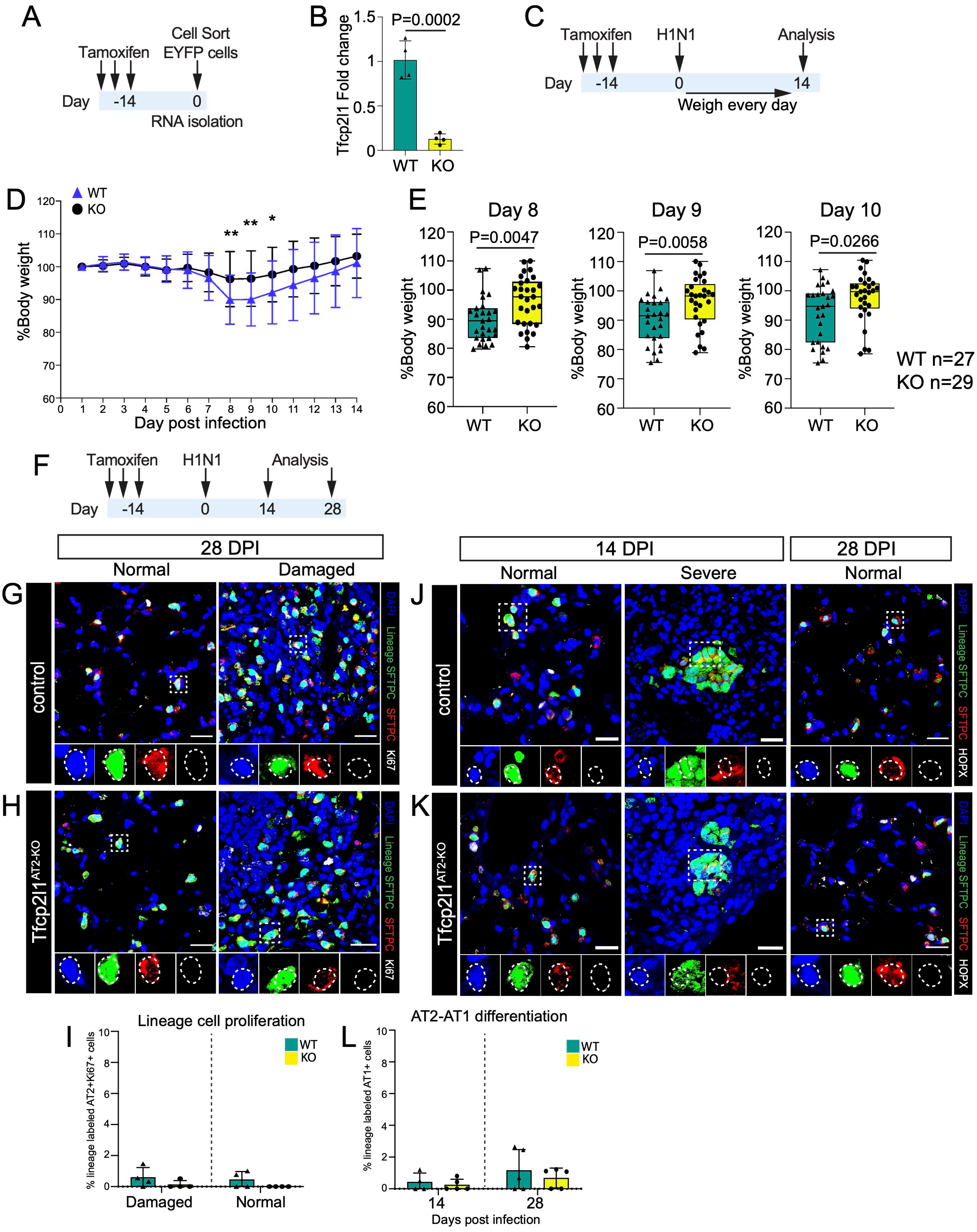

**Supplemental Figure 2. Cardenas et. al.**

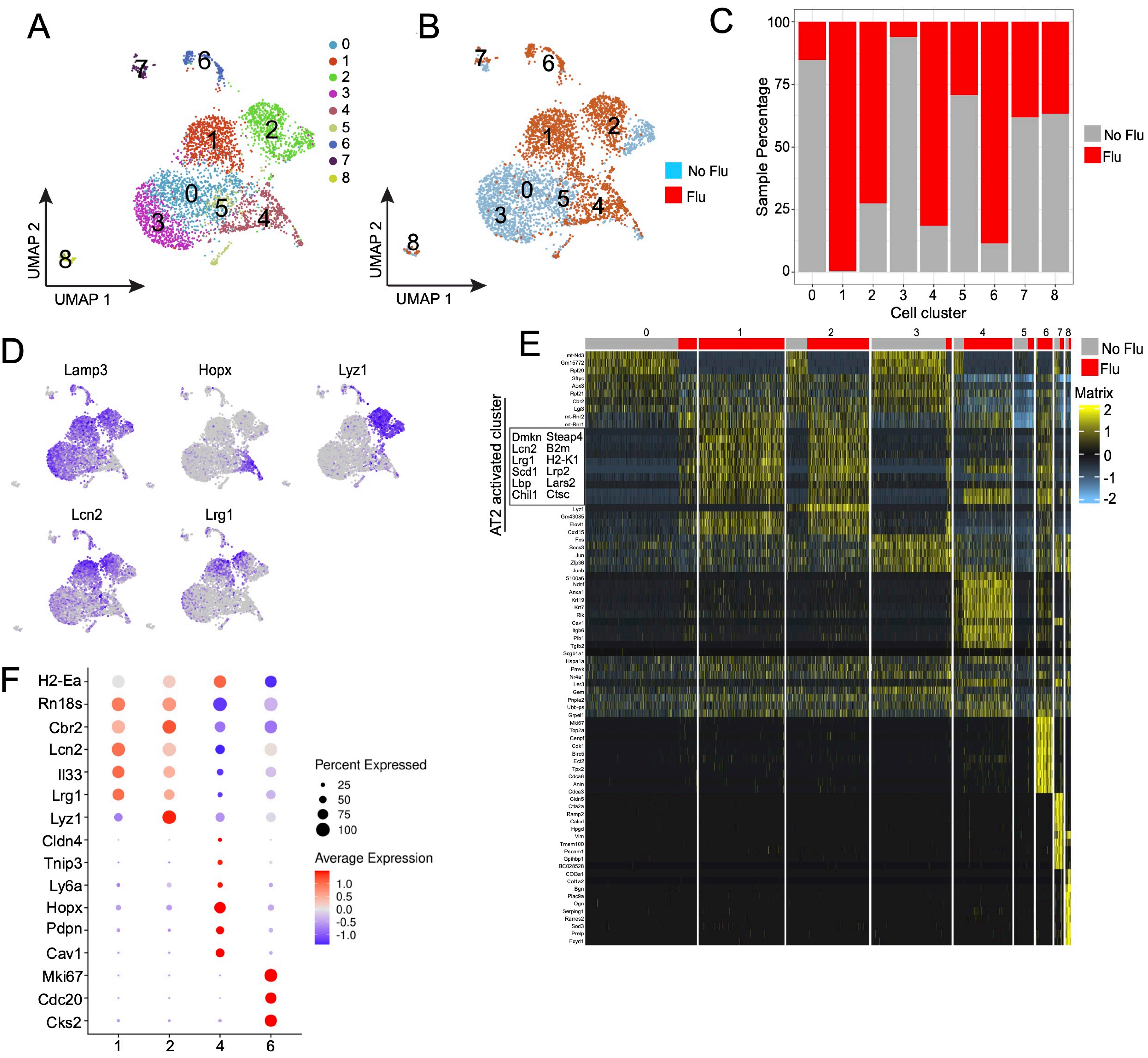


G

| | | <u>adj. P value</u> |
|---|---|---|
| GO:0002474 | antigen processing and presentation of peptide antigen via MHC class I | 6.44506E-07 |
| GO:0019882 | antigen processing and presentation | 1.25214E-06 |
| GO:0048002 | antigen processing and presentation of peptide antigen | 1.78705E-06 |
| GO:0002699 | positive regulation of immune effector process | 1.42079E-05 |
| GO:0002822 | regulation of adaptive immune response | 2.12296E-05 |
| GO:0001913 | T cell mediated cytotoxicity | 2.44583E-05 |
| GO:0002819 | regulation of adaptive immune response | 2.44583E-05 |
| GO:0002456 | T cell mediated immunity | 2.73819E-05 |
| GO:0002824 | positive regulation of adaptive immune response | 2.73819E-05 |
| GO:0002711 | positive regulation of T cell mediated immunity | 2.73819E-05 |
| GO:0002821 | positive regulation of adaptive immune response | 2.73819E-05 |
| GO:0002708 | positive regulation of lymphocyte mediated immunity | 3.11549E-05 |
| GO:0002428 | antigen processing and presentation of peptide antigen via MHC class Ib | 3.11549E-05 |
| GO:0002475 | antigen processing and presentation via MHC class Ib | 4.48627E-05 |
| GO:0002697 | regulation of immune effector process | 4.48627E-05 |

**Supplemental Figure 3. Cardenas et. al.**

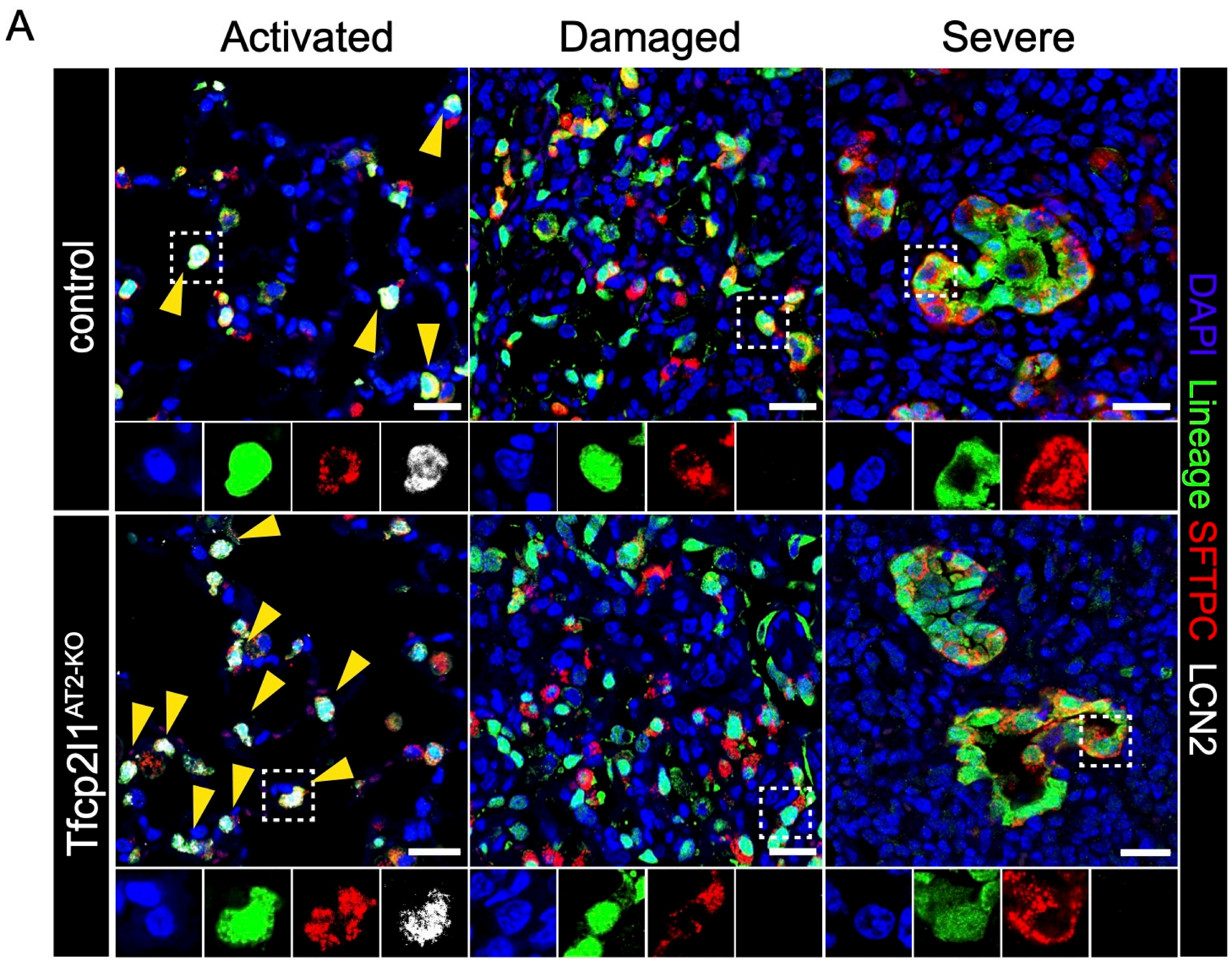

**Supplemental Figure 4. Cardenas et. al.**

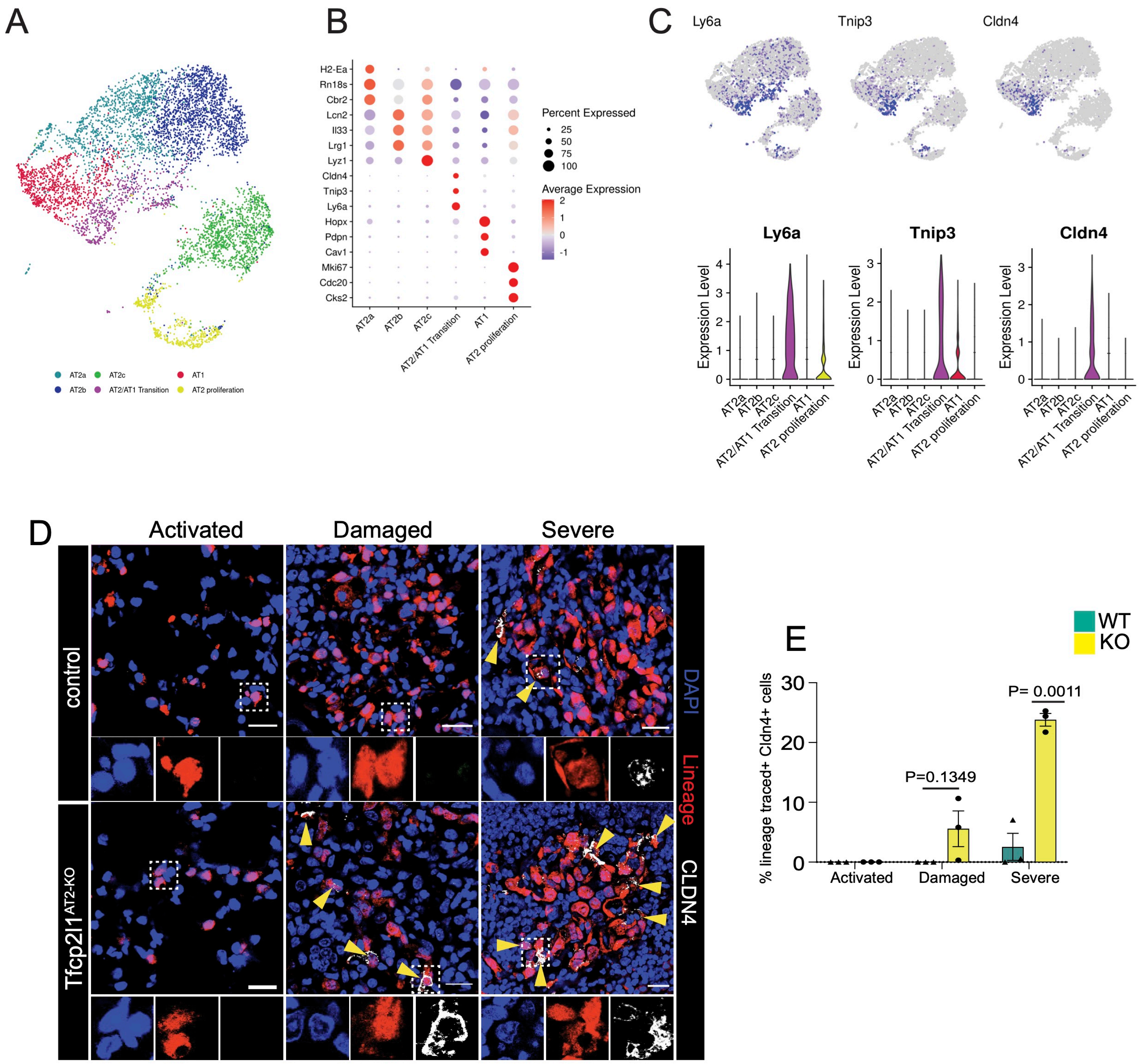

**Supplemental Figure 5. Cardenas et. al.**

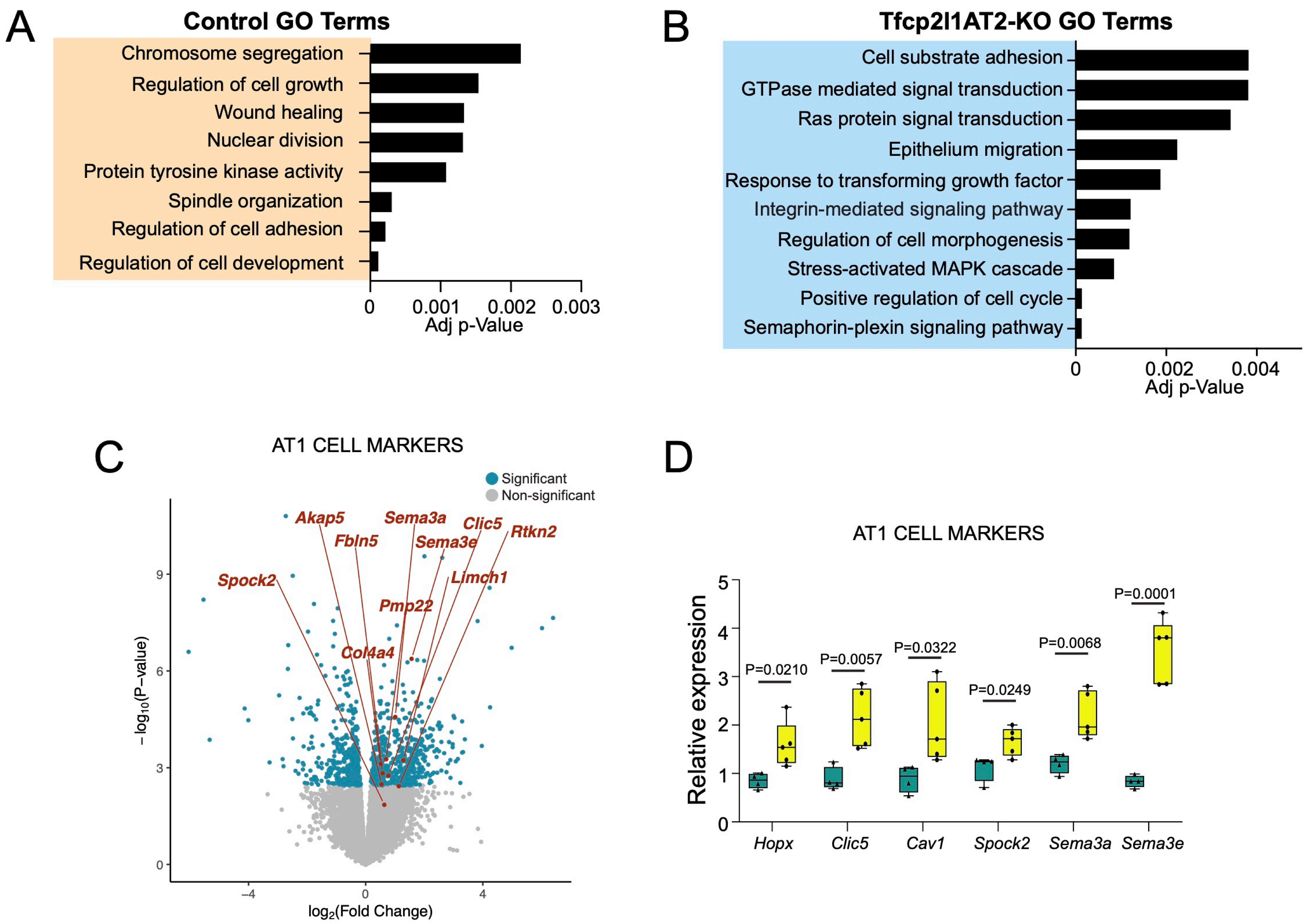
